# Continuous monitoring reveals protective effects of *N*-acetylcysteine amide on an isogenic microphysiological model of the neurovascular unit

**DOI:** 10.1101/2021.03.26.433307

**Authors:** Isabelle Matthiesen, Dimitrios Voulgaris, Polyxeni Nikolakopoulou, Thomas E. Winkler, Anna Herland

## Abstract

Microphysiological systems mimic the in-vivo cellular ensemble and microenvironment with the goal of providing more human-like models for biopharmaceutical research. We report the first such model of the blood-brain barrier (BBB-on-chip) featuring both isogenic human induced pluripotent stem cell (hiPSC)-derived cells and continuous barrier integrity monitoring with <2-minute temporal resolution. We showcase its capabilities in the first microphysiological study of nitrosative stress and antioxidant prophylaxis. Relying on off-stoichiometry thiol-ene epoxy (OSTE+) for fabrication greatly facilitates assembly and sensor integration compared to the prevalent polydimethylsiloxane devices. The integrated cell-substrate endothelial resistance monitoring allows us to capture formation and breakdown of our blood-brain barrier model, consisting of co-cultured hiPSC-derived endothelial-like and astrocyte-like cells. We observe clear cellular disruption when exposing the BBB-on-chip to the nitrosative stressor linsidomine, and report on the barrier permeability and barrier-protective effects of the antioxidant *N*-acetylcysteine amide. Using metabolomic network analysis, we further find drug-induced changes consistent with prior literature regarding, e.g., cysteine and glutathione involvement. A model like ours opens new possibilities for drug screening studies and personalized medicine, relying solely on isogenic human-derived cells and providing high-resolution temporal readouts that can help in pharmacodynamic studies.

## 1. Introduction

The blood-brain-barrier (BBB) maintains the homeostasis of the brain by tightly regulating influx and outflux of everything from metabolites to pathogens and therapeutics. The healthy BBB consists of a layer of brain microvascular endothelial cells (BMECs) closely interlinked by membrane protein complexes.^[1,2]^ This tight barrier is supported structurally and metabolically by pericytes and astrocytes embedded in an underlying layer of extracellular matrix (ECM), and further interacts with nearby neurons and microglia. With its central role in homeostasis, understanding the BBB is critical to shedding light on disorders and developing new therapeutics.

In-vivo studies of the BBB are often complex and unreliable (or even impossible, at least in humans) due to the difficulty in accessing the BBB without impairing its function. Static in-vitro studies recreating the BBB are more accessible and have been performed with several different cell sources, typically on microporous membranes suspended inside standard tissue culture wells (e.g., Transwells).^[1,2]^ More recently, engineering approaches to create <u>dynamic</u> in-vitro models using microfluidics have emerged (i.e., BBB-on-chip).^[3]^ To date, ~20-30 distinct BBB-on-chip models have been reported, attempting to better mimic biological cues such as ECM or fluid flow.^[4,5]^ The most common underlying design is to employ microfluidics cast in poly(dimethylsiloxane) (PDMS), with a porous cell growth support separating two rectangular channels. These approaches have shown great promise, with for instance the application of shear stress-inducing more in-vivo-like morphology in certain BBB cell models, or the cellular compartmentalization shedding light on metabolic coupling between BMECs, astrocytes, pericytes, and neurons.^[6,7]^ An additional advantage of these microengineered systems is the potential for direct integration of sensors to monitor, e.g., neural activity or barrier integrity.^[8]^

In this work, we present a BBB-on-chip – illustrated in **Figure 1a-b** – that builds upon such prior work and combines three crucial advantages: cell ensembles with in-vivo-like functionality, truly continuous barrier integrity monitoring, and a hydrophilic structural material that facilitates simple fabrication and integration of sensors. To date, the combination of two just such advances in barrier-on-chip systems remains elusive.

**Figure 1:**
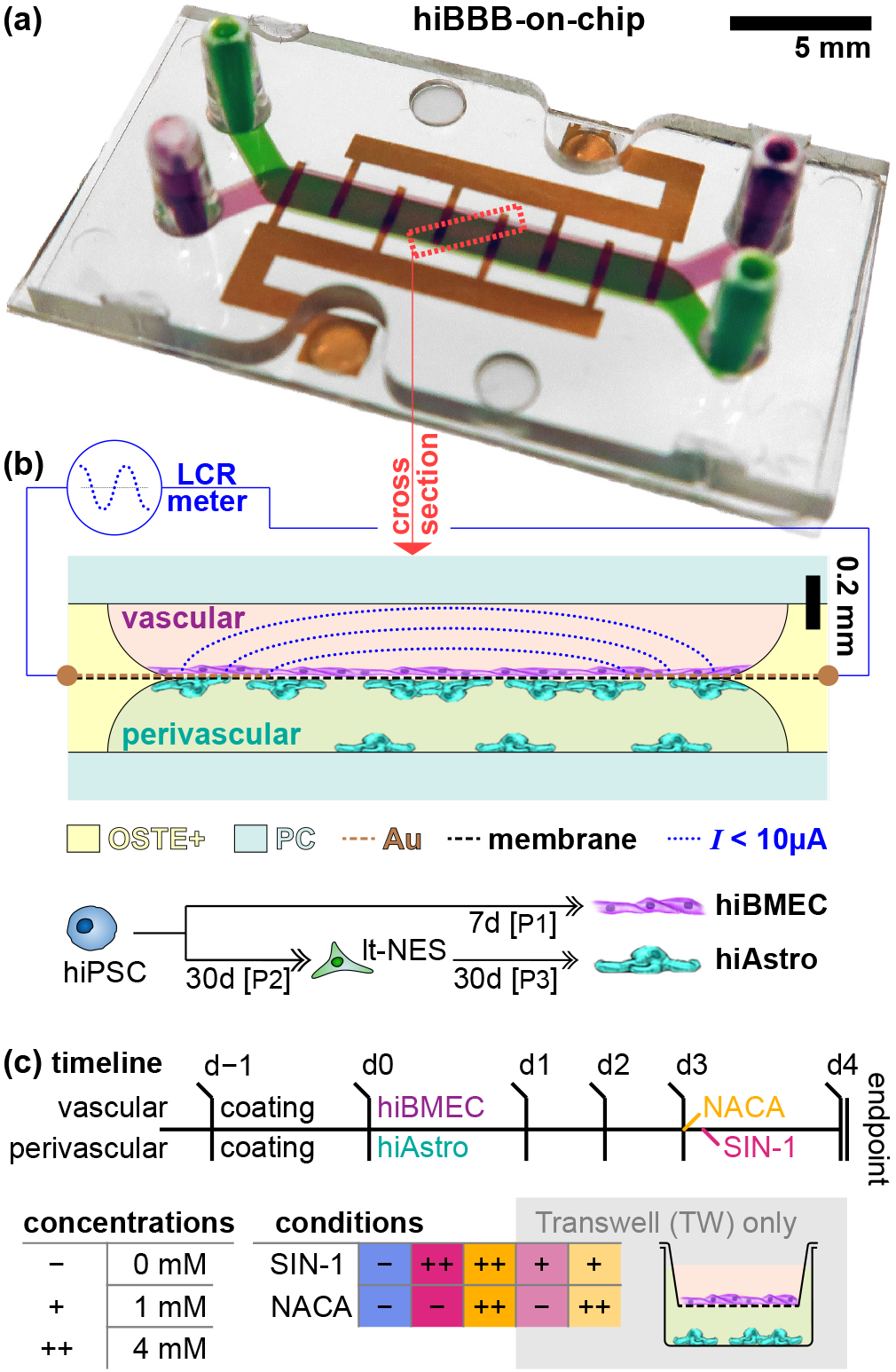
Overview of our microphysiological systems and study design. (a) Photograph of our hiBBB-on-chip system, with the vascular (magenta) and perivascular (green) compartments perfused with colored solutions above and below the permeable membrane with its interdigitated electrodes (gold). The dotted red box indicates the (b) Schematic cross-section of our system, showing the cell co-culture, microfluidic materials, and how the electrodes are used to measure barrier integrity (blue). The various materials utilized are indicated in the legend below and further discussed in the Introduction. P1^[9]^, P2^[10]^, and P3^[11]^ refer to hiPSC differentiation protocols. (c) Experimental timeline and experimental conditions for our study. Abbreviations: off-stoichiometry thiol-ene-epoxy (OSTE+), polycarbonate (PC), gold (Au) human induced pluripotent stem cell (hiPSC), long-term self-renewing neuroepithelial-like stem cells (lt-NES), hiPSC-derived brain microvascular endothelial-like cells (hiBMEC), hiPSC-derived astrocyte-like cells (hiAstro), Linsidomine (SIN-1), *N*-acetylcysteine amide (NACA).

One of the most important functional characteristics in any BBB model is proper integrity of the endothelial layer, where standard assays include permeability of hydrophilic tracer molecules (e.g., fluorescent dyes) and transendothelial electrical resistance (TEER). TEER relies on electrodes placed on either side of the biological barrier to measure the electrical resistance across. In frog and rat BBB, in-vivo TEER has been measured in the range of 1.5– 8 kΩ cm^2^.^[12]^ Conversely, in-vitro culture of primary or immortalized BMEC (human and animal) rarely exceeds 0.5 kΩ cm^2^. Only over the past decade, human induced pluripotent stem cells (hiPSC) have emerged as an alternative to avoid inter-species variation and achieve in-vivo biological function in-vitro.^[1]^ Several protocols have been published for hiPSC-derived BMEC-like cells (hiBMEC) that achieve up to 4 kΩ cm^2^ as monocultures. As mentioned earlier, in-vivo BBB function is reliant also on BMEC interactions with other cell types, with, e.g., tight junction formation being induced by neural progenitor cells.^[2]^ Indeed, co-culture systems with astrocytes and/or pericytes have proven to more fully re-capitulate BBB characteristics from P-gp activity to disease phenotypes.^[1,2]^ For our system, we rely on such hiBMEC,^[9]^ co-cultured with astrocyte-like cells derived from the same hiPSC line (hiAstro; **Figure 1b**). ^[11]^ Such isogenic co-culture can uniquely enable patient-specific disease and treatment models. However, only three other BBB-on-chip studies employ these cellular models (hiBBB);^[13–15]^ most continue to rely on primary or immortalized cells, or just one hiPSC-derived cell type.

TEER is an obvious candidate for sensor integration into BBBs-on-chips, and has been demonstrated in multiple systems, including two of the hiPSC models mentioned above.^[13,14]^ These examples, however, continue to provide measurements only once per day, losing out on a wealth of temporal information. In the aforementioned works (and also non-hiPSC ones like them), this is often due to reliance on manually inserted wire-based (rather than microfabricated) electrodes and associated difficulties in creating long-term incubator-stable interfaces. One prior BBB-on-chip system has demonstrated TEER with minute-scale resolution; however, the study relied on primary cells and recorded only ~12 hours of treatment-phase data.^[16]^ Another study of note, modeling the placenta barrier with a cancer cell line monoculture, has recently employed interdigitated electrodes to monitor cell-substrate endothelial resistance (CSER;^[17]^ another electrical measure of barrier integrity) with subminute resolution.^[18]^ Both these systems, however, suffer from complex fabrication and assembly of 5-9 layers (discussed further in the next paragraph). An important advantage of CSER is a co-planar electrode geometry, which allows for simpler incubator-suitable chip-to-world interfaces. For our hiBBB-on-chip, we similarly implemented CSER with microfabricated interdigitated electrodes in direct contact with the hiBMEC (**Figure 1b**) to gain an electrical measure of hiBBB tightness. We can thus, for the first time, gain barrier integrity information with <2-minute temporal resolution in a hiPSC-based microphysiological system.

Last but not least, extant BBB-on-chip systems are based almost exclusively on PDMS or commercialized plastic devices.^[5]^ The latter are inherently fixed designs not suited to customization with, e.g., sensors. Assembly of PDMS-based microphysiological systems – especially with sensor electrodes involved – on the other hand requires chemical adhesives or physical pressure sealing between the 5+ layers which generally results in long fabrication times and poor reproducibility.^[16,19]^ Less common approaches such as glass (as with the CSER-integrated system mentioned above)^[18]^ or 3D printing suffer from similar sensor integration problems. Many of these methods also necessitate the use of thick layers (PDMS, physical adhesives, gaskets) that put the cells outside the working distance of high-resolution objectives. PDMS in particular is moreover sub-optimal for on-chip cell culture due to its hydrophobicity and strong absorption of small molecules (such as pharmaceuticals).^[20]^ To overcome these challenges in our hiBBB-on-chip, we employ off-stoichiometry thiol-ene-epoxy (OSTE+), a commercially-available polymer system combining the ease-of-fabrication of PDMS, the hydrophilicity and tightly crosslinked (i.e., non-absorbing) nature of thermoplastics, as well as intrinsic bonding capabilities to a range of plastics and metals.^[21]^ It allows us to simplify fabrication to essentially a 3-layer process and to achieve excellent imaging capabilities. OSTE+ has previously been demonstrated for microfluidic cell culture,^[21]^ but never before with multi-day culture of sensitive hiPSC-derived cells.

In this work, we employed our hiBBB-on-chip to study nitrosative stress (i.e., reactive oxygen/nitrogen species, RONS) and its prevention. An excess of RONS, produced, e.g., by microglia in response to cytokines, is a common neuroinflammatory pathway implicated in Alzheimer’s, multiple sclerosis, schizophrenia, and other pathologies.^[22–24]^ Previous studies in traditional cell culture models showed that such RONS stress damages BBB integrity by attacking tight junction proteins ZO-1 and claudins.^[25,26]^ To counteract the effects of nitrosative stress, drugs and antioxidants like *N*-acetylcysteine (NAC) have been studied clinically as treatment, amelioration, or prevention for RONS-related disorders.^[27]^ To date, the biological effects of RONS have not been investigated in barrier-on-chip systems, with the exception of the simplest ROS (H_2_O_2_) as a stressor in primary-or cell line-based endothelial models.^[28–30]^ There are no reports using NAC or any of its derivatives in an on-chip system. Our study is the first to venture into RONS, specifically applying them in the perivascular space where they would be naturally produced by activated M1 (pro-inflammatory) microglia.^[31]^ We rely on the chemical RONS generator linsidomine (SIN-1); it and its prodrug find application both in-vivo and in-vitro.^[32–34]^ SIN-1 generates superoxide and nitric oxide (which further combine to peroxynitrite; thus mimicking microglia activation)^[31,35]^ over the course of a few hours.^[36]^ On the vascular side we applied *N*-acetylcysteine amide (NACA),^[37]^ a more lipophilic (and thus more bioavailable and BBB-permeable) – albeit much less widely used – version of the parent molecule NAC, both proven to reduce neurotoxicity.^[38–40]^

## 2. Results & Discussion

### 2.1. Device Design

Our system, displayed in **Figure 1**, utilized a vertically-stacked channel geometry common in barrier-on-chip systems. A track-etched polycarbonate (PC) membrane – patterned with interdigitated gold electrodes for CSER measurements – served as the cell culture support between the two mirror-symmetric channels. To circumvent separate bonding and alignment of the membrane and the fluidics, we implemented a single-step molding process that first encapsulates the membrane inside (semi-crosslinked) open-channel OSTE+ fluidics. A similar approach has been previously published for PDMS,^[41]^ and we adapted it here for the first time to OSTE+. Unlike the PDMS process, subsequent channel enclosure (with a commercial polycarbonate chip-to-world interface, and a polycarbonate film) was simply facilitated by the epoxy-based bonding properties inherent to semi-crosslinked OSTE+. Besides process simplification, our approach offers two additional advantages. First, it allowed us to produce a unique channel profile with gradual sidewall curvature towards the membrane, which eliminates the typical sharp corners that present likely cell barrier failure points. Second, the top and bottom closures from high-quality thermoplastics mean this is the only material in the optical path. This eliminates scattering and/or autofluorescence issues associated with microfluidic molding processes in general, and previous OSTE-based systems in particular.^[21]^ Additionally, since the bottom polycarbonate film has a thickness akin to a #0 coverglass, imaging of cells on the membrane is possible with even <300 μm working distance objectives (e.g., ~NA 1.2 oil immersion). We note that we attempted further improving the optical qualities by replacing PC with TOPAS or Zeonor; however, these materials proved unable to bond with OSTE+ without further modification (coverglasses, on the other hand, were feasible, albeit yielding fragile devices).

### 2.2. Study Design

We followed the experimental timeline in **Figure 1c** with *N*=23 hiBBB-on-chips, as well as *N*=33 gold-standard Transwell (TW) equivalents, spread out across three (TW: four) independent experimental repeats. Before applying interventions on d3, we verified formation of intact hiBMEC barriers – and excluded compromised ones from subsequent interventions (cf. Section 2.5.1). In brief, we found that intact hiBBBs show permeability measures in line with literature reports. We then applied a prophylactic dose of NACA to randomly-selected intact hiBBBs in the vascular compartments. With a 90-minute delay, we additionally introduced SIN-1 into all of the perivascular compartments. We assessed biological outcomes on d4 (endpoint) using immunocytochemistry (Section 2.3), untargeted metabolomics (Section 2.4), and barrier integrity (Section 2.5). Moreover, we were able to follow the dynamic barrier integrity response using our integrated CSER sensors (Section 2.5.3).

### 2.3. Imaging

In fluorescent imaging, we observed positive immunostaining for the tight junction marker ZO-1, showing a well-delineated network of cell borders in the vascular-compartment hiBMEC (**Figure 2a-b**). The morphology closely resembled our Transwell control group (**Figure 2c**) as well as hiBMEC literature. The one difference we note is that cells are more densely packed in devices. We roughly quantified this in terms of effective (circular) cell radius *r*’ with an interquartile range IQR [4.2; 6.8] μm (versus [5.1; 8.0] μm in Transwells). The analysis additionally reveals hiBMEC in devices to be somewhat less circular, as captured by a compactness ratio (= perimeter / (2 π *r*’)) with IQR [1.8; 3.0] (vs. [1.4; 1.8]).

**Figure 2:**
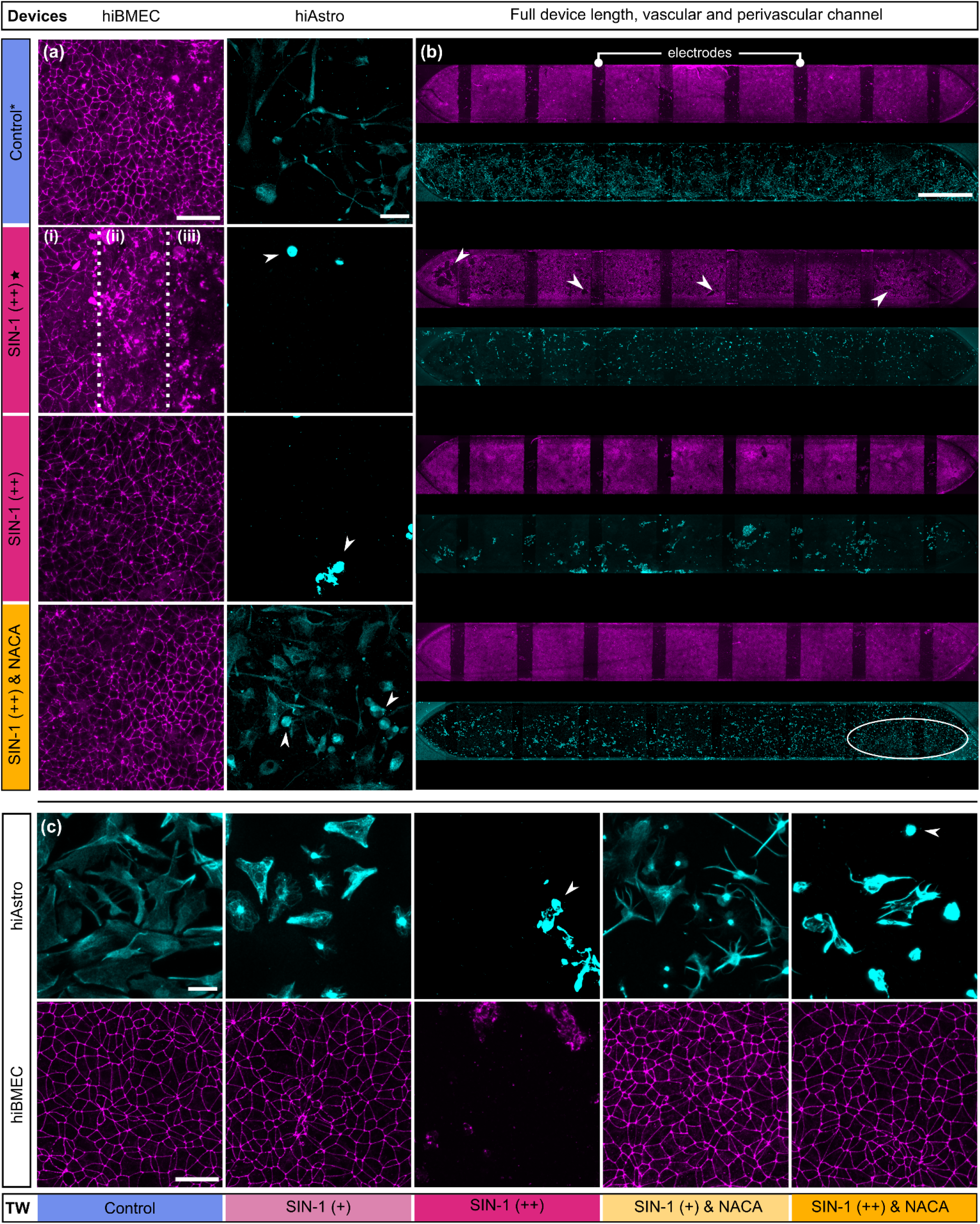
Confocal micrographs of hiBBB-on-chip and Transwell cultures. hiBMEC are stained for tight junction marker ZO-1 (magenta) and hiAstro for astrocytic marker S100B (cyan). (a) High-magnification and (b) corresponding whole-compartment images under our various experimental conditions. Without intervention (control*; see Section 2.5), hiBMEC form a barrier layer with clear tight junctions in the vascular compartment, whereas hiAstro with high S100B expression are distributed across the perivascular channel. SIN-1(++) treatment results in tight junction breakdown and formation of larger defects in approximately half the cases (✶); in the other half, barriers remain intact. The barrier breakdown denoted by zones (a, i–iii) and the white arrows (b) is described in the text. Barriers additionally exposed to NACA (++) remain indistinguishable from controls. Conversely, a majority of hiAstro have detached or condensed (arrows) in devices treated with NACA (++) and/or SIN-1 (++) A small fraction of hiAstro remain attached and spread out, with a comparable morphology to those in the control group (white circled area). (c) Similar trends are seen in Transwells, where hiBMEC exhibit intact barriers in all conditions except SIN-1 (++). hiAstro condense and detach to a large extent when treated with SIN-1 at both 1 mM (+) and even more so at 4 mM (++) doses (white arrows). NACA prophylaxis for lower-dose SIN-1 (+) results in higher hiAstro survival. With a higher dose of 4 mM SIN-1 (++) rescue with NACA is not successful. Scale bars (a) and (c) are 50 μm, (b) 1500 μm.

Our hiAstro, distributed throughout perivascular compartments, exhibited clear expression of astrocytic markers S100B (**Figure 2a-b**) and CD44 (Transwells only; **Figure S1**). Cells were sparser and appear to feature a spindlier morphology in devices than in Transwells, likely due to the presence of media flow (which is less biomimetic, but necessary to study tracer dyes and metabolomics).

#### 2.3.1. Effects of SIN-1 and NACA on hiBMEC

With the application of 4 mM SIN-1 (++), two sub-populations emerged for the hiBMEC: One of these populations – accounting for roughly half of the chips, evenly distributed across the independent experimental replicates – retained fully intact barriers and tight junctions, with indistinguishable morphology from the control* group. In the other half (✶), however, we observed significant barrier disruption. In **Figure 2a** we illustrate the types of disruption across three zones (see also **Figure S2** for nuclear stain): (i) the tight junction network remains intact as seen by clear ZO-1 boundaries between cells; (ii) we observe a partially disrupted network with ZO-1 in disjointed clusters; and (iii) the cell layer itself is disrupted, with voids in the barrier (detached cells) surrounded by cells that are condensed or feature clustered ZO-1 expression. The voids (bordered by the other zones) can be quite extensive and are distributed throughout the devices, as seen in **Figure 2b** (white arrows). Regarding the two apparent subpopulations of hiBMEC response to SIN-1, we refer to further discussion in the context of later Sections (and their additional observations) but note here that we can rule out imaging or fixation artifacts on their basis. Regardless, the prophylactic application of 4 mM NACA (++) proved highly effective in preventing damage to the hiBMEC, with no apparent differences in cellular network morphology compared to the control* group across devices.

The preventative effect of NACA on nitrosative stress damage was wholly replicated in our Transwell hiBMEC (**Figure 2c**). When subjected only to SIN-1, we observed that their barrier integrity was correlated with concentration – at the 4 mM (++) level, no cells remained on the membranes (see also **Figure S2** for nuclei), indicating widespread death and/or loss of adhesion (the lack of even cell debris is likely a wash-related artifact, which are on balance more vigorous compared to laminar wash buffer perfusion in devices). With lower-dose 1 mM (+) exposure, conversely, Transwell hiBMEC barriers showed no differences compared to controls (additional NACA administration also did not elicit further changes).

Our observations of barrier damage caused by SIN-1 (where present), in both the hiBBBs and Transwells, are consistent with literature reports. Severe fragmentation of ZO-1 morphology was documented for peroxynitrite exposure of rat arterial sections.^[42]^ In another study with primary hBMEC, SIN-1 exposure caused a loss of claudins and occludins (though not ZO-1),^[43]^ which would result in ZO-1 becoming unmoored inside the cell cytoplasm, something that our images support (**Figure 2a**). The same study further demonstrated increased matrix metalloproteinase activity leading to ECM degradation, explaining cell detachment. Concentration-wise, existing studies have reported survival at ~70% compared to control, up to 1 mM SIN-1, with significant cell death at >2 mM, in line with our Transwell results.^[44,45]^ The trends we observe are also similar to an hBMEC study showing that 2-hour prophylactic application of 1 mM NACA was successful in reducing or preventing barrier damage from 2.5– 5 mM methamphetamine, an inducer of oxidative stress.^[46]^ NACA prophylaxis against direct application of RONS, as we demonstrated here, had however not previously been demonstrated.

#### 2.3.2. Effects of SIN-1 and NACA on hiAstro

Unlike hiBMEC, hiAstro-on-chip responded to SIN-1 consistently with massive cell death. This is expected due to their more direct exposure (being cultured in the compartment SIN-1 is administered in) and more sensitive nature (as single cells of CNS parenchymal nature, instead of a continuous sheet of vascular-like cells). A differing response may also be linked to inherent differences in cellular uptake, as previously observed between endothelial cells and fibroblasts.^[47]^ The result matches literature reports of primary neurons exhibiting near-complete apoptosis with >12 h exposure to even 1 mM SIN-1.^[35,48]^ One of these studies further showed how SIN-1 exposure results closely matched those achieved via co-culture of activated microglia.^[35]^ Of the hiAstro that remained in the channels, nearly 100% were fully condensed. NACA prophylaxis had a less pronounced effect on hiAstro than on hiBMEC, preventing complete cell death and condensation only for a fraction of cells near the outlets (i.e., where NACA could diffuse across the whole length of the barrier). We illustrate this in **Figure 2b** by circling the area where we identified rescued astrocytes, i.e., those remaining fully attached and spread out. High-magnification images (**Figure 2a**; also **Figure S1**) were taken in these regions to highlight the relevant trends. It appears that, in spite of short diffusion distances across the membrane, vascularly-provided NACA was largely unable to prevent massive perivascular damage from acute RONS exposure.

These findings largely held in the Transwells, with <1% hiAstro survival for SIN-1 (++) (independent of NACA prophylaxis) as estimated from cell coverage (**Figure S3**). We observed that NACA increased low-dose SIN-1 (+) cell survival from 53% (CI_95%_ [27; 81]) to 68% (CI_95%_ [68; 92]). This is in line with a previous study where NACA was proven to counteract glutamate-induced oxidative stress on neuronal cells.^[49]^ The lack of NACA effect for SIN-1 (++) on hiAstro can be attributed to the longer diffusion distance and less favorable compartment mass balance for Transwells compared to devices.

### 2.4. Metabolomics

Untargeted UPLC/MS (ultra-high performance liquid chromatography / mass spectrometry) metabolomics provides a wealth of information about the small (<1 kDa) molecules consumed and excreted by the cells. The approach is by nature broadly exploratory, rather than fully definitive. By separating all our analysis by sampling compartment – vascular, perivascular, and (to account for conditions with disrupted barriers, i.e., non-negligible compartmental mixing) pooled compartments – we can gain insight into their respective contributions. Specifically, we employed two analysis pipelines: one aimed at individual compounds and supported by MS/MS, the other aimed at network-level changes (cf. Methods). Depending on the pipeline, we detected around 2,700 compounds in the supernatant. It is worth noting that these numbers are on the lower end for this type of study (e.g., ~1/3–1/2 compared to our prior work with Caco-2- or primary BBB-on-chip).^[6,50]^ We attribute this to the relatively minimal, protein-poor media we employ here.

#### 2.4.1. Verification of dosage levels

While untargeted metabolomics is not suitable for absolute quantification, we can still obtain information about both NACA and SIN-1 (or, more accurately, its stable breakdown product SIN-1C). We were able to assess SIN-1C as qualitative ratios (**Figure S4**); importantly, we could infer that concentrations in SIN-1 (++) devices were not significantly different between disrupted and intact barriers, and matched (++) media dose controls. This suggests that the observed stochastic behavior of the endothelial layer was not concentration-related. We did find that concentrations in high-dose Transwells were higher than even media controls, possibly due to an increased Transwell evaporation rate altering intended dilution ratios. For NACA, where we could rely on a calibration curve and much better UPLC/MS signal quality, we were able to establish that vascular concentrations in both devices (IQR [3.8; 4.5] mM) and Transwells (IQR [3.8; 4.8] mM) fell within our target range (**Figure S4**).

Based on perivascular concentrations, we further found that NACA successfully permeated all our hiBMEC barriers. We can report the first estimate for its hiBBB *P_app_* = 6 nm s^-1^ (CI_95%_ [4; 8]; fully overlapping for both devices and Transwells). This is broadly in line with our expectations for passive paracellular permeability of a small, hydrophilic molecule (162.2 Da, log*P* ~ —1.5; cf. Section 2.5.1). While the literature does not provide prior examples for NACA, one study with an epithelial Caco-2 barrier model has reported *P*_app_ for NAC below their assay detection limit (~5-10 nm s^-1^).^[51]^ Compared to hiBMEC, Caco-2 are typically leakier (e.g., 13 versus <2.5 nm s^-1^ for sodium fluorescein).^[9,52]^ Taken together, this may indicate that indeed NACA features higher BBB permeability than NAC. However, exemplary drugs with highly effective passive BBB permeability typically rely on a transcellular route and have exhibited *P*_app_ in the range of 100+ nm s^-1^ in a similar Transwell hiBBB model.^[53]^ Our results – including a comparison with our later tracer dye results (Section 2.5.1) – thus suggest that NACA transcellular permeability is present but limited, and the less-efficient paracellular pathway remains relevant if not dominant. While the amide modification may thus indeed improve NACA bioavailability compared to NAC, alternative modification or encapsulation strategies may be beneficial for effective brain delivery.

#### 2.4.2. Principal Component Analysis

We consider first the principal component analyses (PCA), comparing hiBBB-on-chips (and -in-Transwells) for the various treatment conditions. PCA can reveal broad changes in metabolic profiles by considering a wide range of compounds, but without any specificity. The first four PCs account for ~50% of the variation; in **Figure 3a** we show the two PCs with the closest grouping of media controls in each case (see **Figure S5** for the other two PCs, capturing largely within-group variation). Under control conditions – where, unlike with endpoint imaging, we could rely on measurements from intact pre-intervention hiBBB-on-chips – we observe that Transwells and devices are closely co-located for both compartments, yet distinguishable, indicating similar biological function. The clearest intervention-induced difference emerged in the vascular compartment with NACA prophylaxis, which contrasts with our morphological observations showing little difference between that and controls. In the perivascular compartment, we note opposing trends for our systems: Devices clearly reverted toward media-only, which is in line with the hiAstro death and detachment we observe in imaging and the continuous perfusion of fresh media in the devices (opposed to continuous accumulation in Transwells). The pooled-compartment PCA again shows matching trends between systems, with Transwells simply offset toward the right. One notable feature of interest is the inclusion of conditions with SIN-1 (++)-disrupted barriers (⋆ as identified by permeability; cf. Section 2.5.1), which here are indistinguishable from non-disrupted SIN-1 (++) device barriers. This suggests that metabolic function of the surviving cells in either subpopulation remained broadly similar.

**Figure 3:**
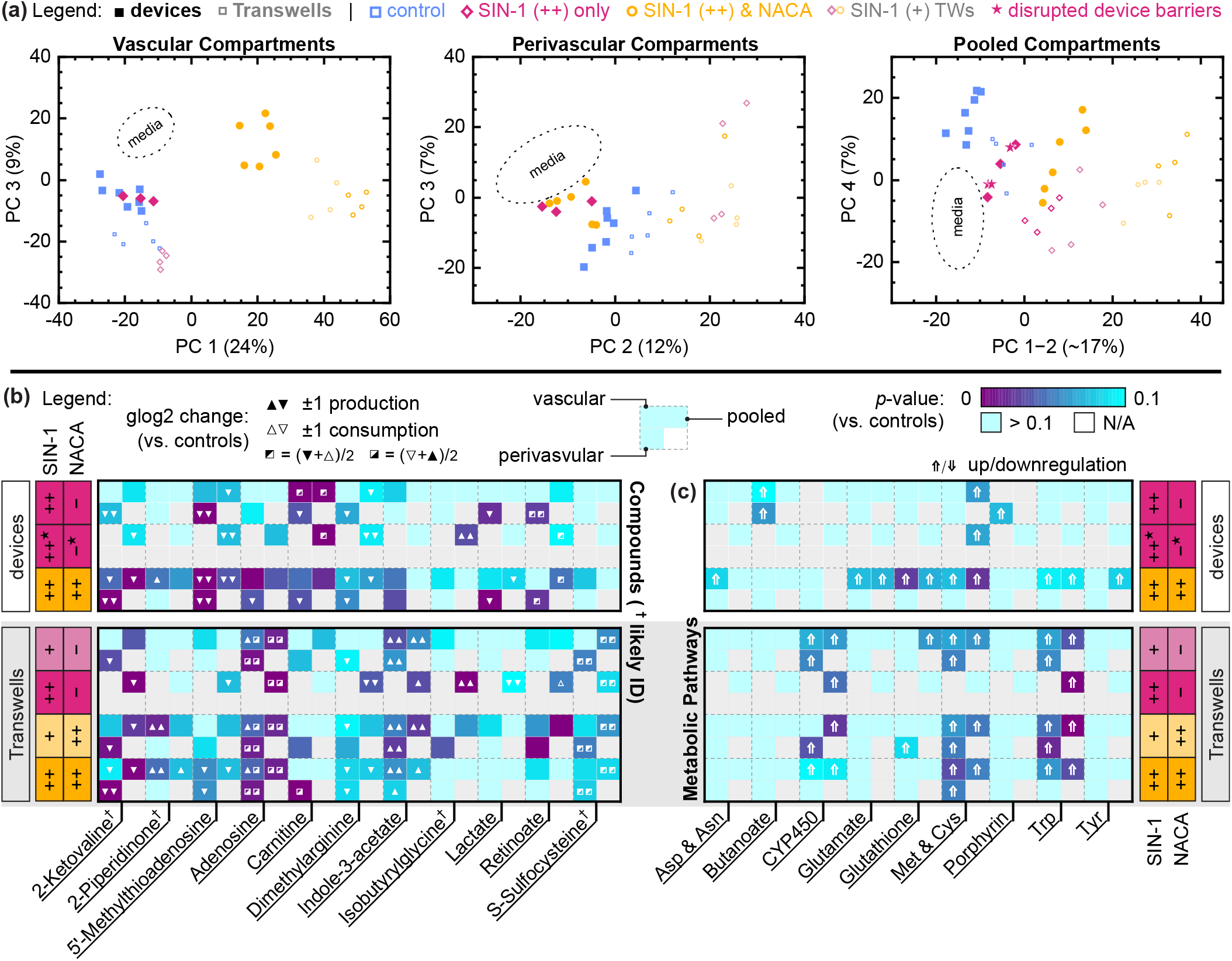
Untargeted metabolomic analysis. (a) Principal Component Analysis (PCA) of untargeted metabolomics, separated by compartment, showing broad changes in metabolic profiles. Symbols indicate devices (filled) and Transwells (small, open), as well as intervention condition. The region containing the majority (80%) of media-only samples is indicated with a dotted outline. PCs (or linear combinations thereof) are selected for least variation in media controls; the remaining 2 (out of first 4) PCs can be found in the Supplement. (b-c) Changes in metabolomic profiles compared to controls on the compound (b) and network (c) level. The matrices are separated vertically by experimental conditions, and horizontally by pathway/compound. The results matrix is very information-dense, but as with all untargeted metabolomic analysis should be considered more for trends than for any single entry. Each cell contains sub-entries on a per-compartment basis (sketch top left), indicating both the respective significance levels (from *p* = 0 (purple) to 0.1 (cyan), or uncolored if not measured). Compound symbols indicate generalized-log2 change in production or consumption (or a change in modality; squares). Network arrows indicate up/downregulation. Compounds were ranked by log (*p* < 0.005)×(|glog2-change| > 1) to select the top 11 endogenous hits. We include all pathways (size > 10) with at least one compartment at *p* < 0.05. Refer to the Supplement for complete lists of potential pathways/compounds.

#### 2.4.3. Networks and Compounds

Here, we consider the putative metabolic changes in the hiBBBs relative to control conditions both for individual compounds as well as at the network level. In **Figure 3b-c**, we show the effective changes as well as significance levels for the top 20. These analyses give higher biological meaning than PCA, but inherently consider a smaller fraction of the underlying data. Overall, our potential pool of hits entails 161 annotated compounds (of which 57 are tentative^†^; cf. **Tables S1-2**) and 24 pathways (**Table S3**). Similar to PCA, we find that NACA prophylaxis induces pronounced metabolomic alterations. On the compound level, we also observe a good share of hits with SIN-1 only. Most of these hits are conserved with NACA prophylaxis; we therefore suggest that NACA does not reverse the metabolic effects of SIN-1. Indeed, all our analyses (PCA, compounds, networks) point to that NACA prophylaxis prevents RONS damage not simply by reverting SIN-1-induced changes back towards controls, but by significant stimulation/regulation of separate metabolic processes. This is in agreement with NAC (the parent molecule of NACA) being only poorly reactive with most RONS (including peroxynitrite, superoxide, and nitric oxide).^[54]^ Its dominant pathway is as a precursor of Cys and glutathione; however, it is believed to affect a wide range of cellular functions through interactions with proteins. Glutathione regulation has also been directly confirmed with NACA in glutamate-stressed neurons.^[49]^

For individual compounds (**Figure 3b**), we differentiate up/downregulation in terms of consumption versus production. The overall patterns we observe in our hiBBBs-on-chips are largely conserved from Transwells, though changes are generally smaller. This is to be expected due to the different mass balance in devices, where continuous perfusion of larger media volumes results in less accumulation. A decrease in production, particularly when present largely in perivascular (or pooled) compartments is likely linked to (particularly hiAstro) cell death, such as for lactate. We ascribe that hypothesis also to 2-Ketovaline, which is a valine metabolite readily produced and released by astrocytes,^[55]^ as well as retinoate, which is produced by astrocytes and metabolized by BMEC.^[56]^ 5’-Methylthioadenosine, on the other hand, shows downregulation also vascularly, more strongly so in the presence of NACA. This may be a result of this metabolite’s involvement in antioxidant and particularly Met pathways.^[57]^ 2-Piperidinone, one of the few consistently upregulated compounds (including devices), does not allow for much analysis due to its nature as a small fragmentary metabolite.^[58]^ Strong upregulation of adenosine production by astrocytes is associated with inflammatory events, and is aimed at modulating inflammatory response.^[59]^ This would be consistent with our Transwell results, where the high levels of adenosine can also pass into the vascular compartment over the experimental duration. Lowered production/increased consumption of carnitine across compartments and conditions in the hiBBBs-on-chips (ameliorated by NACA) is consistent with its function as an endogenous antioxidant (thus reactive with peroxynitrite).^[60]^ The lack of equivalent Transwell response may point to this being a late-stage response, thus more easily picked up in devices. Dimethylarginine is an in-vivo marker of elevated oxidative stress, in apparent contradiction to the downregulation we observe.^[61]^ This likely indicates a more systemic nature of this metabolite’s RONS regulation, which our systems do not recapitulate. Indole-3-acetate is typically associated with the gut microbiome (possibly indicating a false ID), but it can also occur as part of the endogenous Trp metabolism, and may thus correlate with the Trp network results (see below).^[58]^ Isobutyrylglycine and S-Sulfocysteine, meanwhile, are metabolically not well understood;^[58]^ their potential involvement in RONS may warrant further study.

The individual compound results are numerous, with some suggestive of interesting future research. To gain an additional view that is both broader than compounds, but more specific than PCA, we finally turn to network-level analysis in **Figure 3c**. Here, one of the strongest and most well-conserved hits with SIN-1 (conserved with additional NACA) was Met & Cys metabolism. This aligns well with the central role of sulfur-containing amino acids in both pro- and antioxidant interactions.^[62,63]^ Met & Cys are also highly favored targets for direct reactions with peroxynitrite.^[24]^ A third directly-implicated amino acid is Tyr, but this does not appear to extend to pathway-level changes here. NACA prophylaxis additionally revealed a strong hit – particularly in our hiBBB-on-chips – on the downstream glutathione pathway, matching in-vivo findings for the central mechanism behind NAC action.^[54]^ This is further supported by the hit on the closely-linked glutamate pathway. Moreover, we find that Trp is closely linked with NACA application in both devices and Transwells (though in the latter it is also linked to SIN-1, albeit with roughly one order of magnitude lower significance than for NACA). We note one extensive in-vivo metabolomic study of NAC (using lymphocytes from lupus patients) implicates three pathways – Trp, glutathione, and a mitochondrial pathway too small to be included in our analysis.^[64]^ The presence of significant CYP450 regulation – indicated in a range of brain disorders – in Transwells only may reflect the higher survival of hiAstro in that system (and lack of fresh media perfusion even after cell death).^[65]^ Due to the discrepancy also in vascular-side CYP450 response, involvement of this pathway may additionally be early-onset and/or time-limited (which for devices implies significant dilution). For devices, we found additional hits in the Asp & Asn, butanoate, porphyrin, and Tyr pathways. Being each present in only one compartment and condition at *p* < 0.05 calls for more targeted studies to confirm their involvement.

### 2.5. Barrier integrity

#### 2.5.1. Pre/post-intervention (tracer dye)

As mentioned earlier, we relied on d2–3 tracer dye permeability (**Figure 4c**) to confirm formation of *n*=13 intact barriers in our hiBBB-on-chip (an additional *n*=6 will constitute the control* group; Section 2.5.2). Specifically, we employed the small and highly hydrophilic Cascade Blue hydrazide (527.47 Da, log*P* ~ — 7),^[66]^ and established a threshold value of 10 nm s^-1^ (based on the first minimum in our data distribution). The higher device failure compared to Transwells (based on d3 TEER; **Figure 4a**) is to be expected due to continuous flow removing detaching cells. The intact-barrier control 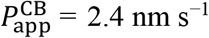 (CI_95%_ [1.6; 3.4]) closely resembles the range of the only other hiBBB-on-chip system where comparable data is available (2–6 nm s^-1^; lucifer yellow, 457.3 Da, log*P* ~ —5).^[15]^ Our observations also fall within the 1–5 nm s^-1^ range of broader hiBBB literature observations (though many of these utilize fluorescein, which is smaller but also less hydrophilic, thus a less optimal paracellular marker; 332.3 Da, log*P* ~ 0).^[1,67]^

**Figure 4:**
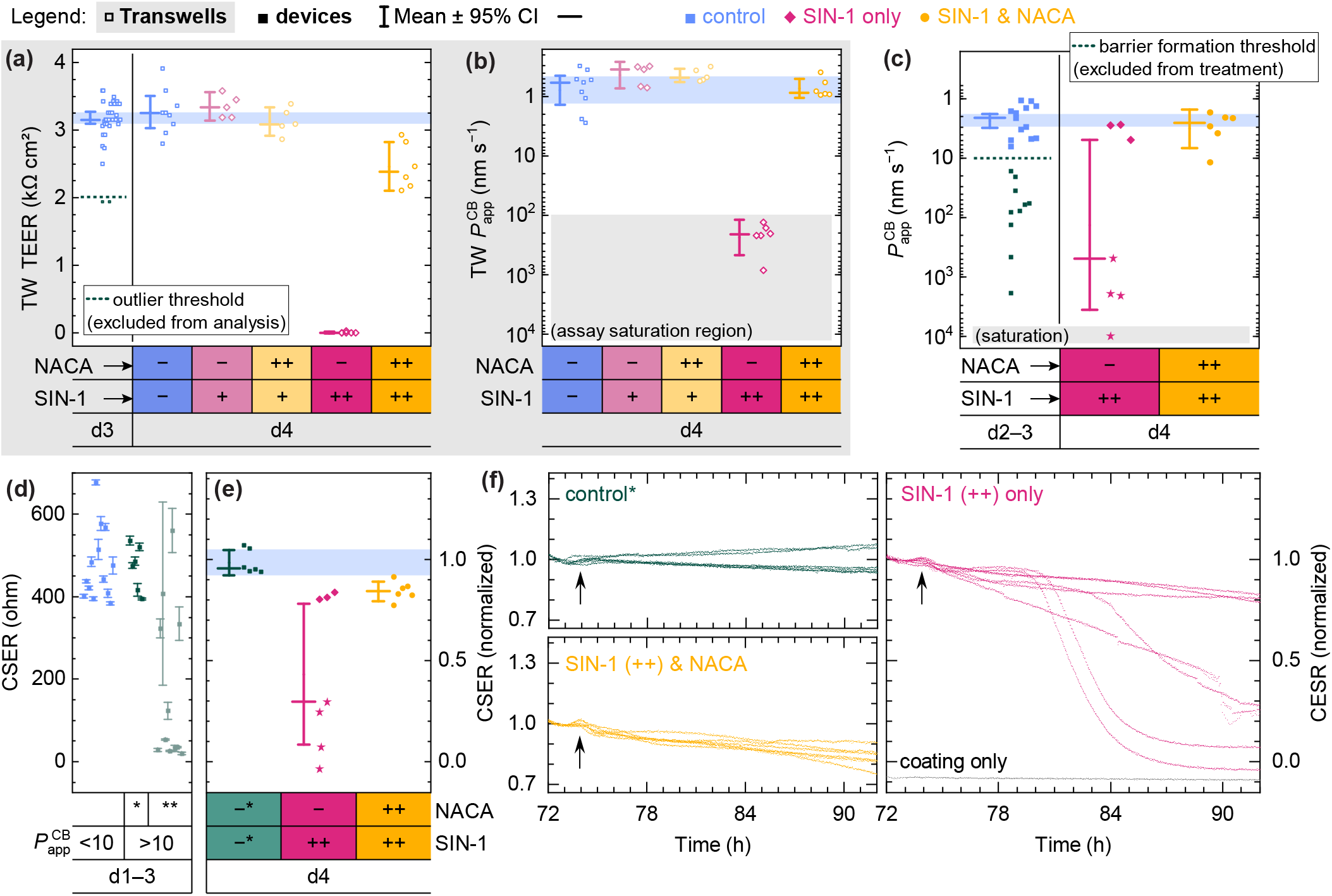
Barrier permeability. Data are separated by condition as per the legend. (a-b) Transwell TEER and tracer dye permeability. (c) Device tracer dye permeability. (d) Preintervention CSER averaged over the signal plateau phase (cf. Figure S6). Data in this graph only are displayed as median ± IQR, and grouped by their corresponding measured 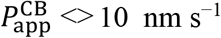. The */** sub-grouping is assigned based on a threshold for the median (100 Ω) and the relative variability (IQR / median; 10%). (e) CSER endpoints for TEER comparison. To highlight intervention-induced changes, the data are normalized to respective pre-intervention signals. (f) Time-resolved normalized CSER, separated by condition.

With our own Transwell controls, we determined 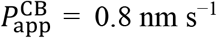 (CI_95%_ [0.4; 1.4]; **Figure 4b**). Some of the discrepancy compared to devices may arise from conservative estimates of cell area and flow rates in the latter. However, we posit that a large part of this discrepancy is related to our microscopy observations of the hiBMEC shape (cf. Section 2.3). Specifically, the smaller cell area and longer cell perimeter add up to a significantly (from our parameter estimates, ~1.9-fold) longer tight junction network in the devices, thus allowing more opportunities for paracellular permeation.^[68]^ We further note that the higher device 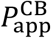 may be considered more in-vivo-like, where fluorescein permeability has been measured at 14.6 nm s^-1^ in live rat brain capillaries.^[69]^ Interestingly, the significant system-based discrepancy here does not match our earlier observation of identical NACA permeability in devices and Transwells. For the antioxidant, this substantiates a contribution also of a passive transcellular route (which would remain unaffected by tight junction network length).

When we applied 4 mM SIN-1 (++) perivascularly to the intact hiBBBs-on-chips, 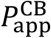 jumped to 130 nm s^-1^ (CI_95%_ [4.9; 3500]) by d4 (**Figure 4c**). The data exhibit a clear bimodal distribution, with *n*=3 remaining practically unchanged compared to controls, whereas *n*=4 (✶) show clear barrier breakdown with 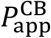 in excess of 100 nm s^-1^. We reiterate that each of our three fully-independent experimental replicates contained at least one of either scenario. The intact/disrupted barrier sub-populations here correlate with those from imaging. The on-chip hiBMEC response does, in a sense, closely match the Transwells: one sub-population similar to SIN-1 (++) Transwells (> 100 nm s^-1^), the other – like lower-dose SIN-1 (+) Transwells – matching the pre/no-intervention controls. However, the response appears stochastic rather than concentration-dependent (discussed further in Section 2.5.3). Importantly, we observe that an additional prophylactic application of 4 mM NACA (++) to *n*=6 devices was wholly effective in preventing significant barrier breakdown, with 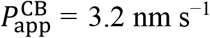 (CI_95%_ [1.5; 6.6]) largely overlapping the control confidence interval.

#### 2.5.2. Pre/post-intervention (electrical)

For the CSER data from our integrated sensors, we first consider summarized results akin to those obtained with 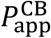 (and thus similar to daily Transwell TEER). Through d3, the intact barriers (*n*=13) plateaued at 490 Ω (CI_95%_ [440; 530]; **Figure 4d**; see also **Figure S6**). The relative confidence interval here is wider than in Transwell TEER (**Figure 4a**), which we attribute to fabrication process variability. Specifically, we suspect that the variable (and difficult-to-control) metallization depth inside the pores is a factor. Still, we find that a drop of CSER signals below 100 Ω, or signal fluctuations (measured as IQR / median) in excess of 10%, always corresponded with a visibly disrupted barrier (**). To highlight this, for **Figure 4d** only, we include also *N*’=7 devices (all within one experimental repeat, otherwise excluded) where erroneous cell seeding resulted in limited barrier formation, followed by breakdown/delamination within 24–48 h (confirmed by visual brightfield inspection). However, due to the locally-sampled nature and somewhat higher inherent variability of CSER, confirmation of complete continuity by tracer dyes is required (**Figure 4c**). Those devices with stable CSER but above-threshold 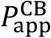 (*; *n*=6) – i.e., featuring barrier defects away from the electrodes, often not apparent even on visual inspection as in **Figure 2b** – were retained as the control* group, considered only for imaging and CSER.

To reduce the influence of device variability when considering the impact of SIN-1 on barrier integrity, we normalized the CSER data for the intervention period (d3–4) with respect to the per-device values recorded immediately prior. The resulting endpoint data (**Figure 4e**) closely mirror those from 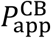 (**Figure 4c**), with the control* group remaining unchanged at 0.99 (95% CI: 0.92–1.05), while SIN-1 (++) caused a drop to 0.43 (95% CI: 0.08–0.78) by d4 – with a bimodal signal distribution matching the tracer dye assay. Unlike with 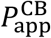, however, prophylactic NACA (++) did not fully protect measured barrier integrity, showing instead a small but significant decrease to 0.84 (95% CI: 0.79–0.89). Interestingly, this mismatch between CSER and 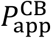 is very similar to that between TEER and 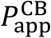 for Transwells (**Figures 4a-b**). The measurable change in electrical, but not (tracer dye-) molecular permeability compared to no-intervention controls suggests that the induced RONS damage in the NACA prophylaxis group is largely confined to very small (ion-size) defects. This is akin to a prior study showing some antioxidants fully preventing RONS damage to primary hBMEC in terms of *P*_app_, but not TEER.^[43]^ Overall, not only do our CSER sensor results reflect independently-measured alterations in barrier permeability; their trends also align with gold-standard methods. Based on data from both the pre- and post-treatment phase, we can further calculate that in spite of the aforementioned limitations our sensors achieve 71% sensitivity, and notably 100% specificity, in barrier breakdown detection.

#### 2.5.3. Dynamic measurements

As mentioned in the Introduction, hiBBB-on-chip studies to date show at best daily sensor readouts similar to what we present above. The advantage of our CSER sensors here is their continuous measurement capability with <2-minute temporal resolution. We want to briefly mention the barrier formation phase (**Figure S6**), where we find that the barrier formation time course is different from TEER, with CSER leveling off from d1 (rather than d2 for TEER). In part, we attribute this to performing CSER using a permeable membrane (including partially-metallized pores), where a conduction path through the perivascular channel is always available. CSER signals in general are moreover impacted not only by tight junction formation, but also cell attachment and spreading. Our selection of a more resistive-dominated (rather than ionic double layer-dominated) measurement frequency at 6 kHz seeks to maximize the tight junction contribution, and is in line with typical CSER frequencies for such experiments.^[17]^ Based on the time course, it is also clear that mere cell attachment – happening within 1–3 h – is not the dominant factor. However, cell spreading likely contributes to our early signal plateaus. Importantly, our observations match some^[70,71]^ (though not all)^[72]^ prior studies of CSER with hiBMEC (using monoculture in on commercial, non-permeable substrates). The inconsistencies in literature are likely due to a strong dependence of CSER on the precise electrode geometry.^[17]^ The daily impedance measurement time course presented by Motallebnejad with their on-chip hiPSC co-culture model also shows a similar pattern to ours, with a plateau from d1.^[14]^

Of larger interest for our present study are the full measurement traces during the intervention period shown in **Figure 4f** (normalized as in 2.5.2). 72 h marks the 4 mM NACA (++) administration into the vascular compartments of our hiBBBs-on-chips (where applicable); timing of the 4 mM SIN-1 (++) in the perivascular compartments, with a ~1.5h delay, is indicated by the arrows. Up to that point, we observe a small down-up oscillation across conditions (including control*), likely reflecting cell response to freshly-prepared media. Control* barriers (blue) remain very flat after this point – most of them decreasing to ~0.95 by the endpoint, with two slightly increasing. This may be a result of small defects from d3 (which this group is known to include based on *P*_app_) being present on top of the electrodes for these two devices, and the hiBMEC gradually closing them up over the course of d4. Already by 75h, we note a distinct drop in CSER signals to ~0.95 for all SIN-1 (++) devices compared to that control* group. Differences from NACA prophylaxis (yellow) only arise around 76h, with those barriers only showing a very gradual decline to the aforementioned 0.84 endpoint signals. The CSER data from SIN-1-only devices (magenta), on the other hand, exhibit more varied characteristics. Three devices remain practically indistinguishable from those that also received NACA (matching 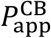 data). The other four devices show CSER dropping below 0.2, but with unexpectedly varied time courses – from largely continuous (~20 h) degradation to more sudden (~4–6 h) loss of integrity, with onset times varying from 75 h to 82 h. Besides the bimodal nature of barrier breakdown already seen in 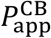, the CSER data thus further reveals stochastic dynamics of the breakdown process.

As we have previously stated, the equal distribution of intact and disrupted SIN-1 (++) barriers across our three independent experimental replicates indicates these dynamics are not an artifact arising out of any batch effects (e.g., cellular differentiation, device coatings, etc.). Our metabolomics data moreover suggest that SIN-1 (++) concentrations were largely similar between the devices in question (cf. Section 2.4.1), and no clear sub-group distinction regarding SIN-1barrier breakdown (✶) appeared in PCA (cf. **Figure 3a** and **Figure S5**). Based on all the available evidence, we posit a process whereby SIN-1 (as it approaches a certain threshold concentration) causes minor barrier damage which cannot be wholly prevented by NACA. These initial defects are likely ion-sized, based on the decrease in electrical signals (CSER/TEER; based on ionic currents) while the larger-molecule 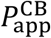 remains constant. However, NACA prevents any larger-scale damage. We hypothesize that major hiBMEC disruption happens when the small defects accumulate across the barrier, allowing for *random* formation of clustered defects where larger-molecule diffusion can occur. This would result in SIN-1 penetration into the vascular compartment, where it can more directly disrupt the hiBMEC across the entire area and, in combination with fluid shear stress, accumulate further damage leading to cascade barrier failure. The concentration of SIN-1 in our devices is not sufficiently above the threshold concentration, however, to guarantee that clusters of defects are consistently formed, leading to bimodal observations. In Transwells, on the other hand, SIN-1 (++) is sufficiently above the threshold concentration to ensure breakdown, due to the significantly higher mole-per-area (based on a much larger perivascular compartment) as well as potentially higher liquid evaporation (cf. Section 2.4.1.). Lower-dose SIN-1 (+), meanwhile, remains below the threshold. We have, however, recorded heterogenous SIN-1 response in prior Transwell experiments using TEER, where the SIN-1 concentration was closer to the threshold (**Figure S7**; not directly comparable due to different media).

While verification of the biological mechanisms underlying heterogeneity will require further study, this does not diminish the added value provided by continuous CSER monitoring in our hiBBBs-on-chips. It allowed us to follow the temporal dynamics of barrier alterations (or lack thereof), depending on the applied interventions, with unprecedented temporal resolution, yielding a rich new dimension to hiBBB studies. Such temporal resolution is especially important for acute biological injuries such as stroke or traumatic brain injury, as well as to study the pharmacokinetics and pharmacodynamics of potential treatments, where 15–30-minute timescales are highly relevant.^[73–75]^

## 3. Conclusion

We presented a new type of hiBBB-on-chip system based on OSTE+ microfluidics, which enabled simple integration of both the permeable membrane as well as sensors to monitor barrier function continuously. Our isogenic hiBBB co-culture in this microfluidic system exhibited properties similar to both traditional cell culture models and to the few existing hiBBB-on-chip systems. Specifically, we observed formation of continuous tight junction networks and expression of the glial marker S100B in the respective cell types, and verified junction integrity using fluorescent tracer diffusion. When applying perivascular nitrosative stress – akin to microglia activation – using the peroxynitrite generator linsidomine (SIN-1), we observed consistent and widespread hiAstro cell death, and hiBMEC barrier disruption in approximately 50% of cases. Prophylactic vascular administration of the antioxidant NACA increased hiAstro survival fractionally. However, we found that its hiBBB permeability was low compared to drugs classified as BBB-permeable. On the vascular side, NACA proved wholly effective in preventing barrier disruption. The network-level metabolomic response profile we observed moreover aligned with the available literature findings for nitrosative stress and NAC, particularly stimulation of the glutathione pathways in the cells that help prevent damage caused by RONS exposure. Last but certainly not least, we were able to monitor changes in barrier integrity using the integrated CSER sensors with (for hiBBB systems) unprecedented temporal resolution. Our study thus revealed for the first time the effects of direct RONS application on the BBB, as well as the effects of NACA prophylaxis, in a microphysiological system.

We recognize that, compared to TEER, our CSER-based sensor approach samples the barrier integrity locally rather than globally, thus requiring confirmation of the intact-barrier state from a global measure such as tracer dye permeability. Drops in CSER, notably, were 100% specific with regard to barrier disruption. We also expect that optimization of the fabrication process will allow us to lower signal variability between devices in the future. Overall, we believe the advantages of a co-planar electrode layout in terms of fabrication complexity and off-chip interfacing (a critical factor in facilitating continuous measurements) outweigh the caveats. In terms of biological functionality, we acknowledge that a lack of pericytes in our hiBBB co-culture presents a limiting factor. However, markers for pericyte lineage are significantly less well-established than those for endothelial cells or astrocytes.^[1]^ Research on this front is rapidly evolving, however, and we expect to utilize a hiAstro-hiPeri mixture for the perivascular compartment in the near future. Another goal is to introduce microglia to such BBB model where onset of neuroinflammation could be biologically rather than chemically induced. Such a model could help understanding cell-cell interactions in the BBB, e.g. gain insight on the dual effect of microglia on BBB integrity – namely their neuroprotective effect during inflammatory states but also microglia-astrocytic endfeet interactions for example causing barrier disruption.

In conclusion, we have demonstrated a hiPSC-based microphysiological system of the BBB featuring truly continuous monitoring, with RONS barrier disruption and antioxidant prophylaxis as a proof-of-concept. We have validated that OSTE+ is not only suitable for simple fabrication of – and sensor integration in – microphysiological systems, but also for long-term culture of sensitive hiPSC-derived cells and high-quality imaging. Our hiBBB-on-chip can find immediate utility in the study and screening of drugs with barrier-modulating effects, providing unique readouts about temporal pharmacodynamic profiles. Based as it is on isogenic co-culture, it is furthermore well-suited to disorder-specific studies, such as shedding light on the potential causative involvement of BBB leakage in neurodegenerative and neuropsychiatric disorders.

## 4. Experimental Section

Detailed methods can be found in the Supplemental Material.

In brief, we fabricated OSTE+ devices in a double-molding process that directly integrates track-etched PC cell growth supports. Starting form a CNC-machined positive aluminum master mold, we created negative PDMS-based molds that were clamped around the PC membrane for reaction-injection molding of OSTE+.^[76]^ Prior to integration, we patterned interdigitated gold electrodes for measuring CSER onto the membranes.^[77]^ The resulting open-channel OSTE+ structures on either side of the membrane were sealed with polycarbonate to form the cell culture microfluidics, and disinfected before use.

On d–1, we placed devices into an incubator alongside a peristaltic pump and media reservoirs, and connected the electrodes (via a matrix switching unit) to an LCR meter. We coated the vascular and perivascular compartments with fibronectin/collagen and laminin, respectively. To recreate the vascular hiBBB we thawed and seeded pre-differentiated hiBMEC on d0.^[9]^ To recapitulate the astrocyte-endothelial interaction we concurrently thawed and seeded pre-differentiated hiAstro on the perivascular side of the membrane.^[11]^ After cell seeding, we continuously perfused media and measured CSER until d3. On d3, we also collected control-group media for tracer dye measurements and metabolomics (the latter immediately frozen until their off-site analysis). Subsequently, utilizing the continuous media perfusion, we began administering NACA to the vascular channels, to serve as a neuroprotective agent, before introducing SIN-1 in the perivascular channel with a 1.5-hour delay. On d4, we again collected media for permeability and metabolomics. Immediately afterward, we fixated the cells and stained for ZO-1 apically and S100B basally; nuclei were counter-stained with Hoechst. In parallel, we kept Transwell cocultures for comparison of a dynamic vs static model, assaying TEER daily and collecting media for tracer dye permeability and metabolomics on d4. In **Table S4**, we summarize the number of hiBBBs across all our experimental conditions.

Our image analysis focused on maximum intensity projections of the confocal stacks acquired. Besides visual inspection, we further quantified cell coverage (Transwell hiAstro) and cell size (hiBMEC) in ImageJ. Tracer dye fluorescence measurements were used, with the aid of calibration curves, to calculate apparent permeability. These data, together with TEER and CSER, were analyzed in OriginPro. We report values in terms of means and their 95% confidence intervals, excepting cases where the distribution width is of greater importance (in which case we report medians and interquartile ranges). For metabolomics, we followed two parallel workflows, aimed either at annotated compounds using ThermoFisher’s proprietary software,^[78]^ or at network-level results using open-source tools.^[50]^ Since both SIN-1 and NACA acted not only on the cells, but also reacted with the media itself, we employed media background-subtraction in our workflows for more meaningful comparisons. For metabolomics data only, we conducted statistical analyses in R and (by design) needed to rely on *p*-values.

## Supporting Information

Supporting Information is available online or from the author.

## Acknowledgements

<u>T.E.W. and A.H. contributed equally to this work</u>. We are grateful to Mercene AB and SPIBER AB for providing us with OSTE+ and Biosilk, respectively. We thank Violetta Nikiforova for her assistance with hiBMEC differentiation. We appreciate the support of Dr. Charles Vidoudez and the Harvard Center for Mass Spectrometry in the metabolomic analysis, and acknowledge the Biomedicum Imaging Core at Karolinska Institute. We further appreciate the support of Mikael Bergqvist, Cecilia Aronsson, as well as Electrum Laboratory in device fabrication. T.E.W. is grateful for funding from the European Union’s Horizon 2020 Research and Innovation Program under the Marie Sklodowska-Curie grant agreement “NeuroVU” (No. 797777). A.H. acknowledges funding from the Knut and Alice Wallenberg Foundation (No. 2015-0178), Göran Gustafsson Stiftelse, and Forska utan Djurförsök.

## Author contributions

**Isabelle Matthiesen:** Conceptualization (supporting); Methodology (equal); Investigation (equal); Analysis & Visualization (equal); Writing (equal). **Dimitrios Voulgaris & Polyxeni Nikolakopoulou:** Methodology (supporting); Review & Editing (supporting). **Thomas E. Winkler:** Conceptualization (equal); Methodology (equal); Investigation (equal); Software; Analysis & Visualization (equal); Writing (equal); Supervision & Administration (supporting). **Anna Herland:** Conceptualization (equal); Methodology (supporting); Review & Editing (supporting); Supervision & Administration (lead).

