## Supporting Information for "Continuous monitoring reveals protective effects of *N*-acetylcysteine amide on an isogenic microphysiological model of the neurovascular unit"

*Dr. P. Nikolakopoulou, Prof. A. Herland*
*AIMES, Center for Integrated Medical and Engineering Science, Department of Neuroscience, Karolinska Institute,*
*Solnavägen 9 / B8, 171 65 Solna, Sweden*

NB: The supplemental Detailed Experimental Section is located after the supplemental Figures & Tables referred to in the main manuscript.

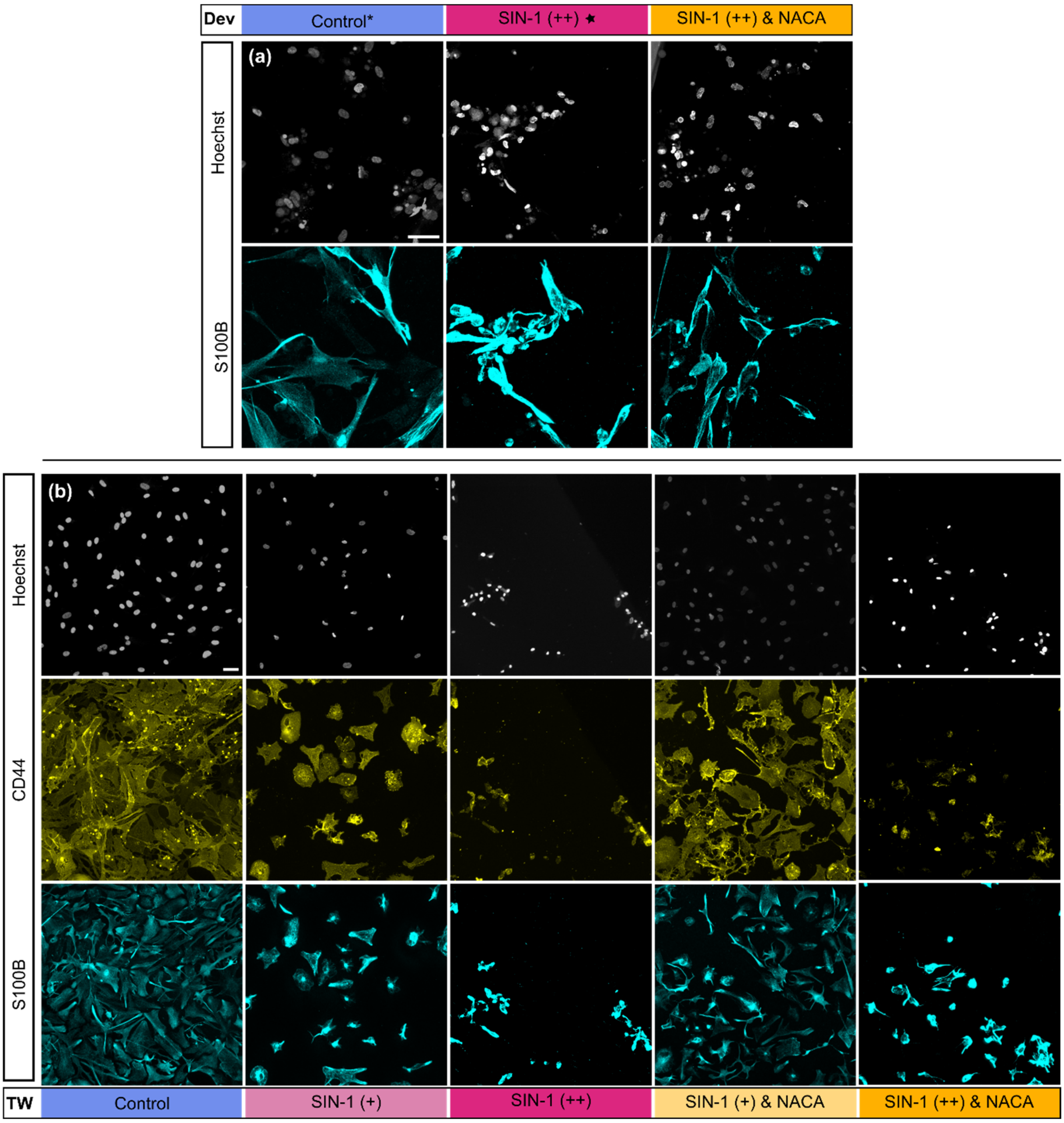


**Figure S1: Confocal micrographs of hiAstro.** In devices **(a)**, hiAstro without intervention (Control) show high expression of astrocytic marker S100B (cyan) and the cells are attached and display a typical astrocytic morphology with spread out extrusions. Moreover, the nuclei (gray) appear rounded without excessive condensed morphology. During treatment with SIN-1 (++) without (and, to a lesser degree, also with) NACA, hiAstro have a more condensed morphology, curling up and detaching (S100B) and nuclei are to a larger extent condensed. In Transwells **(b)** hiAstro without intervention (Control) appear attached and spread out in their morphology (CD44, yellow) and (S100B, cyan), and nuclei (gray) appear rounded. When exposed to SIN-1 (+), hiAstro show less coverage with signs of condensed soma (CD44 & S100B), as well as some condensed nuclei. During NACA prophylaxis (SIN-1 (+) & NACA), hiAstro detach to a lesser extent and have a more spread-out morphology closer to that of the Control. Similarly, nuclei appear rounded as seen in Control. For higher-dose SIN-1 (++), with or without NACA, most cells have detached, and remaining cells appear rounded up and condensed (CD44 & S100B), nuclei also appear mostly condensed. Scale bars are 50 µm.

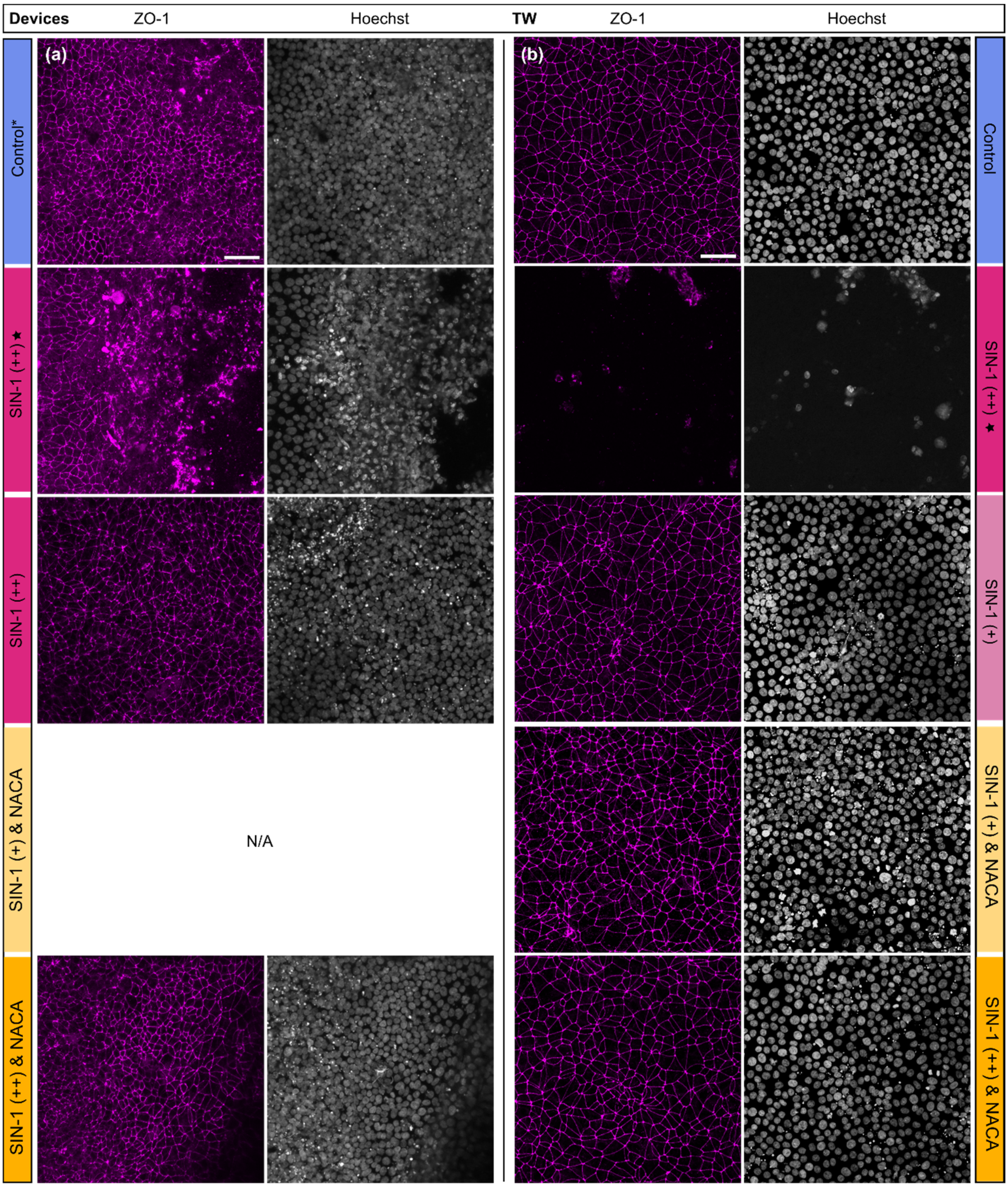


**Figure S2: Confocal micrographs of hiBMEC.** hiBMEC are stained for tight junction marker ZO-1 (magenta) and nuclear stain Hoechst (grey) under our various experimental conditions. In devices **(a)** without intervention (control*; see Section 2.5), hiBMEC form a barrier layer with clear tight junctions and rounded nuclei in the vascular compartment. SIN-1 (++) results in tight junction breakdown and formation of larger voids (no ZO-1 or nuclei) in approximately half the cases (★); in the other half, barriers remain intact. Barriers additionally exposed to NACA (++) remain indistinguishable from controls. **(b)** Similar trends are seen in Transwells, where hiBMEC exhibit intact barriers in all conditions except SIN-1 (++), where we largely observe neither tight junctions nor nuclei. Compared to controls, we find some nuclear condensation at SIN-1 (+) independent of NACA, as well as for SIN-1 (++) & NACA. N/A: condition not applicable. Scale bars are 50 µm.

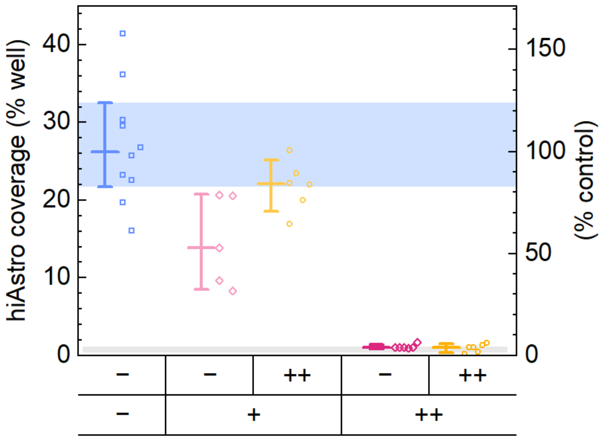


**Figure S3: Surface coverage.** Quantification of hiAstro surface coverage in Transwells, depending on experimental conditions. The shaded gray region represents the region where automated quantification often failed (based on insufficient cell/background contrast), in which case we assigned a conservative estimate of 1(% well) based on visual comparisons with successfully quantified low-coverage wells. Bars represent means ± 95% confidence interval.

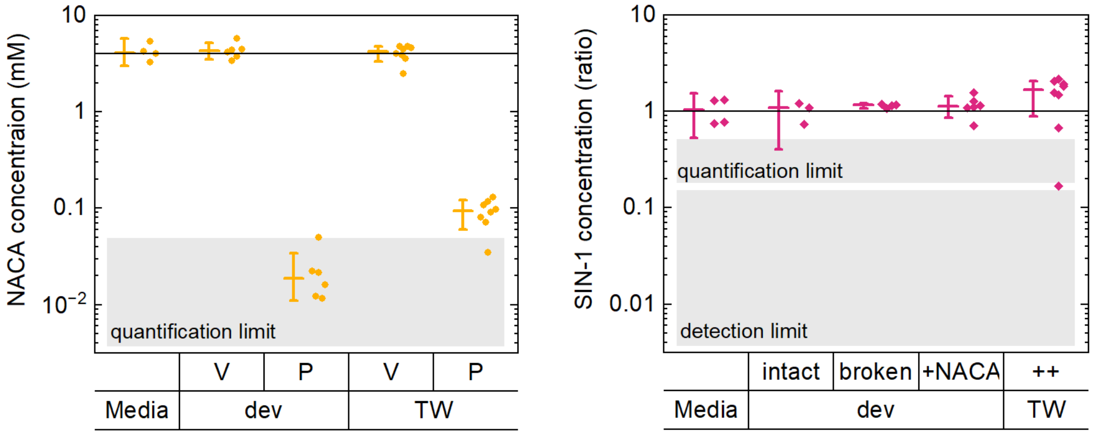


**Figure S4: Concentration analysis. (left)** Semi-quantitative analysis of NACA based on metabolomics, separated by experimental condition, including the vascular (V) and perivascular (P) compartments of hiBBBs-on-chips (dev) and -in-Transwells (TW). All vascular values are clustered close to the target dosage of 4 mM. Perivascular values, while less accurate, allow us to estimate compound permeability. The quantification limit is estimated from noise levels. Bars represent medians ± 95% confidence interval of the mean. **(right)** Qualitative analysis of the SIN-1 breakdown product SIN-1C based on metabolomics. Due to significantly noisier MS signals (compared to NACA), combined with a lack of calibration samples, we assess relative ratios compared to media controls (left). Comparing the three device conditions, we find they fall into a range overlapping the controls. Importantly, between broken and intact barriers, only one sample reveals a somewhat lower concentration – an effect that can thus at most explain one of the three intact barriers. The comparison to Transwells shows generally higher concentrations. One potential explanation is Transwell media evaporation altering the intended dilution ratios when adding SIN-1. The Transwell outlier below the quantification limit may be due to peak misidentification from retention time shifts.

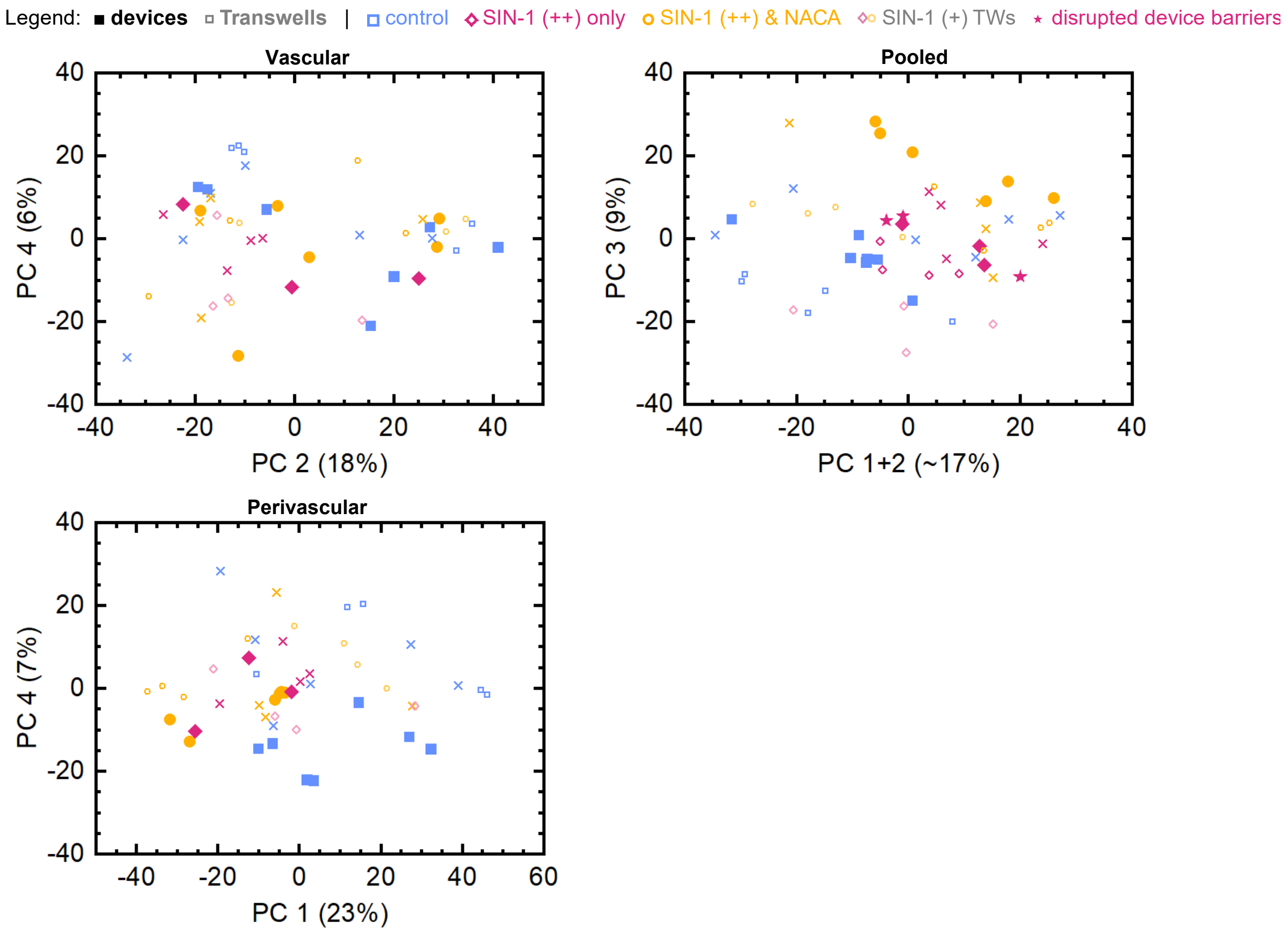


**Figure S5: Principal Component Analysis (PCA)** of untargeted metabolomics, separated by compartment. Symbols indicate devices (filled) and Transwells (small, open), as well as the intervention condition. The two (out of first four) PCs not shown in the main text are displayed here, capturing largely within-group variation, as evidenced by the wide spread of media samples ×.

**Table S1.** Table of annotated compounds discovered in our samples. Annotation is based on MS/MS analysis. **Bold** compounds match the cutoff of $p<0.005$ and net expression change |NES|>1 in at least one compartment. For inclusion in the main manuscript, compounds across both Table S1 and S2 were ranked by log($p$)*|NES|, and non-endogenous compounds discarded.

| Compounds | | | |
|---|---|---|---|
| **Adenosine** | **Hypoxanthine** | Pyruvic acid | Tryptophan |
| Alanine | **Indole-3-acetic acid** | Quinoline | Tryptophan |
| Arachidonic acid | **Indole-3-lactic acid** | Retinoic acid | Tyrosine |
| Arginine | **Lactic acid** | Serine | Uracil |
| Asparagine | Lauric acid | SIN-1C | Valeric acid |
| Asparate | Leucine | Stearic acid | Valine |
| **Betaine** | **Linoleic acid** | Tetramethylpyrazine | 1-Vinylimidazole |
| Bis(methyl-benzylidene)sorbitol | Lipoic acid | Thiamine | 2-(acetylamino)-3-(1H-indol-3-yl)propanoic acid |
| Capric acid | Lysine | Triethyleneglycol dimethyl ether | 2-Amino-4-methylpyrimidine |
| **Caprolactam** | Malic acid | Triisopropanolamine | 2-Hydroxy glutaric acid |
| Caprylic acid | Mesaconic acid | Tris(2-butoxyethyl) phosphate | **2-Mercaptoethanol** |
| **Carnitine** | Methionine | Tryptophan | **2-Methyl-4-isothiazolin-3-one** |
| Choline | Methyl salicylate | Tryptophan | 3-(3,4-dihydroxyphenyl)propanoic acid |
| Cystine | Methylquinoline | Tyrosine | **3-Aminoquinoline** |
| **Dimethylarginine** | Myristic acid | Uracil | 3-hydroxy-6-methoxy-2-phenyl-4H-chromen-4-one |
| Dipropylene glycol | Myristyl sulfate | Valeric acid | 4-(anilinomethylidene)-3-methyl-4,5-dihydroisoxazol-5-one |
| dimethyl ether | N,N-Diethylethanolamine | Valine | **4-Acetamidobenzoic acid** |
| Dodecyl sulfate | N,N-Dimethylaniline | Pyroglutamic acid | 4-Ethynylaniline |
| Erucamide | NACA | Pyruvic acid | 4-Guanidinobutyric acid |
| Ethyl myristate | **Nicotinamide** | Quinoline | 4-Hydroxybenzaldehyde |
| Furoic acid | Nicotinic acid | **Retinoic acid** | 4-Indolecarbaldehyde |
| Glutamic acid | Nonanoic acid | Serine | **4'-Methoxychalcone** |
| Glutamine | **Oleic acid** | SIN-1C | 4-Piperidinecarboxamide |
| Glycerophospho-choline | Ornithine | Stearic acid | 5-Hydroxyindole-3-acetic acid |
| Glycine | **Oxoproline** | Tetramethylpyrazine | 5-Methylcytosine |
| Guanine | Palmitoleic acid | Thiamine | **5'-Methylthioadenosine** |
| Histidine | Pentadecanoic acid | Triethyleneglycol dimethyl ether | 6,6-Dimethyl-4-[2-(2-thienyl)ethyl]-1,2,4-oxadiazinane-3,5-dione |
| Homoserine | Phenylalanine | Triisopropanolamine | 8Z,11Z,14Z-Eicosatrienoic acid |
| Hydroxybutyric acid | P-Hydroxybenzaldehyde | Tris(2-butoxyethyl) phosphate | |
| Hydroxy-D-proline | Prolinamide | | |
| | Proline | | |
| | Pyridoxine | | |

**Table S2.** Table of tentatively (high-likelihood) annotated compounds discovered in our samples, based on partial MS/MS analysis matches. **Bold** compounds match the cutoff of $p<0.005$ and net expression change |NES|>1 in at least one compartment. For inclusion in the main manuscript, compounds across both Table S1 and S2 were ranked by log($p$)*|NES|, and non-endogenous compounds discarded.

| Tentative compounds | |
|---|---|
| **Acamprosate†** | Prolylglycine† |
| Acetylcysteine† | Pyridoxamine† |
| Alanine or sarcosine† | SIN-1† |
| **Alpha-Ketoisovaleric acid†** | Tylosin† |
| Benzotriazole† | Undecanoic acid† |
| Crotonic acid† | Urocanic acid or aminonicotinic acid† |
| Crotonic acid† | **Urocanic acid†** |
| Cystathionine† | Valpromide† |
| Cystathionine† | 1-(2-methoxy-4-nitrophenyl)pyrrolidine† |
| **Cysteine-s-sulfate†** | **1-(3-Amino-2,3-dideoxypentofuranosyl)-5-methyl-2,4(1H,3H)-pyrimidinedione†** |
| Glucose† | **1-(3-Amino-2,3-dideoxypentofuranosyl)-5-methyl-2,4(1H,3H)-pyrimidinedione†** |
| Glucose† | 1,5-Naphthalenediamine† |
| Hymexazol† | 1,5-Naphthalenediamine† |
| **Isobutyrylglycine†** | 1-Tetradecylamine† |
| **Levofloxacin†** | 2,6-di-tert-Butylphenol† |
| Lyxose† | **2-cyano-N-(2-nitrophenyl)benzenesulfonamide†** |
| Methylmalonic acid† | **2-methyl-5-(propylsulfonyl)pyrimidin-4-amine†** |
| N,N-Dimethylglycine or GABA† | 2-Methylhippuric acid† |
| N1-[5-(3,5-dimethylpiperidino)-4-fluoro-2-nitrophenyl]acetamide† | **2-Piperidinone†** |
| **N4-[(dimethylamino) methylidene]-3,5-dimethylisoxazole-4-sulfonamide†** | **4-Acetamidobutanoic acid†** |
| N-Acetylasparagine† | 4-methyl-5,6,7,8-tetrahydro[1,2,4]triazolo[5,1-b]quinazolin-9(4H)-one† |
| N-Acetylputrescine† | 4-Methyl-5-thiazoleethanol† |
| N-Benzylformamide† | 4-oxo-retinoic acid† |
| N-Ethylglycine† | 5-[2-(3-Furyl)ethyl]-8a-(hydroxymethyl)-5,6-dimethyl-3,4,4a,5,6,7,8,8a-octahydro-1-naphthalenecarboxylic acid† |
| Norleucine† | 5-Nitro-o-toluidine† |
| Ofloxacin impurity A† | **7-(2,3-Dihydroxypropyl)theophylline†** |
| **Ofloxacin†** | **[4-(1,2,4-Oxadiazol-5-ylmethyl)-2-morpholinyl]acetic acid†** |
| o-Toluidine† | |
| Pipecolinic acid or nipecotic acid† | |
| Piperonylonitrile† | |

**Table S3.** Pathways from the MFN metabolomic model for which our samples have coverage of at least 10 putative compounds (parentheses), as per mummichog. Smaller pathways are excluded due to the very high likelihood of false positives. Pathways with at least one $p < 0.05$ are **bold**.

| MFN Pathway (# coverage) | | |
|---|---|---|
| Ala & Asp (27) | **Glutathione (17)** | **Porphyrin (12)** |
| Arg & Pro (43) | Gly, Ser, Ala, Thr (39) | Purine (27) |
| Ascorbate & Aldarate (13) | Glycerophospholipid (15) | Pyrimidine (42) |
| **Asp & Arg (67)** | His (20) | **Trp (43)** |
| Beta-Alanine (19) | Lys (18) | **Tyr (59)** |
| **Butanoate (25)** | **Met & Cys (33)** | Urea cycle (56) |
| **CYP450 (12)** | Nicotinate & Nicotinamide (15) | Val, Leu, Ile (20) |
| **Glutamate (19)** | Nitrogen (12) | Xenobiotics (13) |

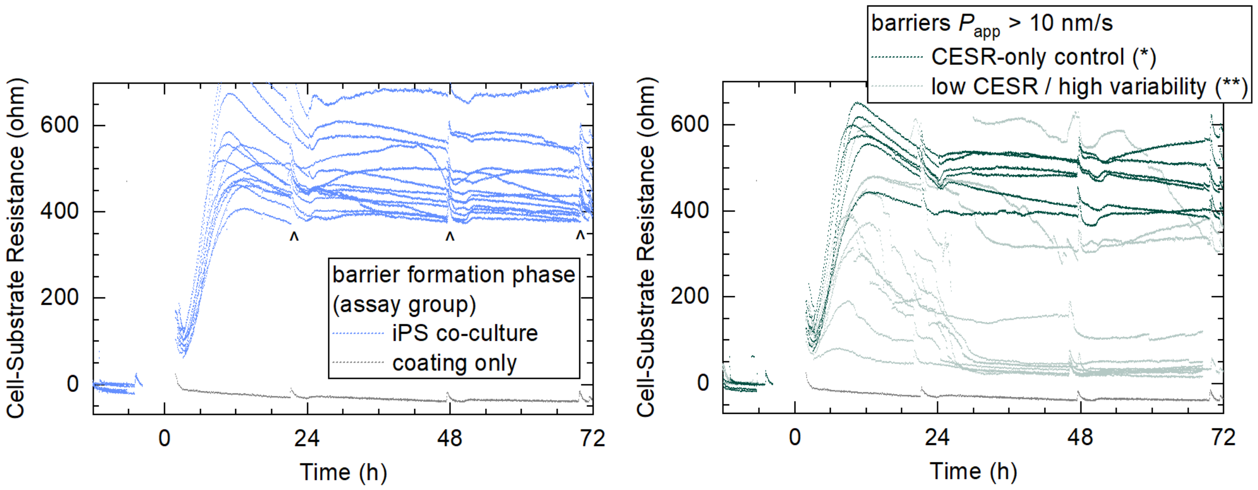


**Figure S6: CESR data from the barrier formation phase. (left)** hiBBBs-on-chips in the assay group, i.e. those showing $P_{app}^{CB}$ < 10 nm/s and subsequently challenged with SIN-1 (and NACA). ^ indicates where media reservoirs were replenished. **(right)** Devices that were classified as disrupted prior to d3 based on > 10 nm/s permeability (or, for *N*'=7, visual inspection). The control* group is indistinguishable from the assay group (left), suggesting small defects that either remained unchanged through the plateau phase or that are located between the electrodes.

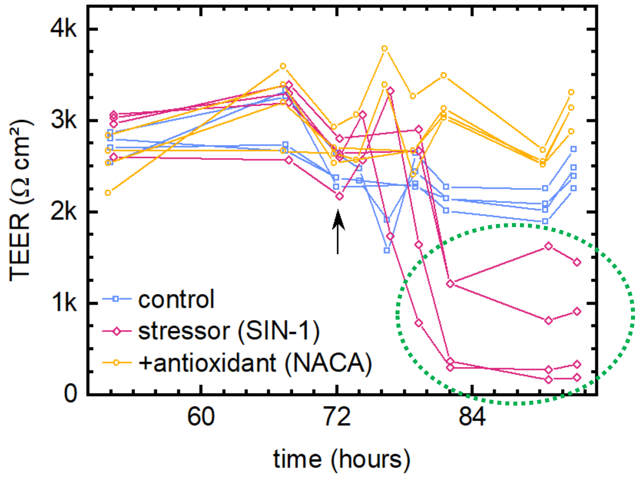


**Figure S7: Heterogeneity in Transwell SIN-1 response.** Transwell TEER data from a prior experiment utilizing media supplementation with 1× B27 (minus antioxidants) instead of ITS, and nominal 2 mM dosages of SIN-1 (supplied by Sigma-Aldrich instead of Biotium) and NACA. The timecourse roughly aligns the one from our present study, with NACA prophylaxis indicated by the arrow, and SIN-1 introduced with a ~2-hour delay. Repeated (rather than just endpoint) TEER measurements in these data – especially between 72-84 h – yielded higher temporal resolution, but could also result in additional stress, and thus lower TEER, for the cell layers. We note that SIN-1 response here was significantly more heterogenous (dotted green circle) than in our present study high-dose Transwells. Although none of the SIN-1 treated Transwells remained fully intact, these data indicate the conditions here may have been similarly close to a SIN-1 threshold as the devices in our present study.

**Table 1.** Number of devices and Transwells considered in this study across the various experimental groups. For certain groups, only a random sub-set was sent for metabolomic analysis, indicated by [brackets]. Threshold exclusion denotes those hiBBBs that were completely excluded from further analysis based on d3 assays. Numbers were evenly distributed across three (devices) or four (Transwells) independent experimental repeats. The only exception are d1–2 devices (§), which represent a separate (fourth) independent repeat.

| | **d1-2 only§** | **d3 (control)** | **threshold exclusion** | **d4 (control)** | **d4 (control*)** | **SIN-1 (++) only** | **SIN-1 (++) & NACA (++)** | **SIN-1 (+) only** | **SIN-1 (+) & NACA (++)** |
|---|---|---|---|---|---|---|---|---|---|
| **hiBBB-on-chip** | 7§ | 23 [7] | −4 | | 6 [0] | 7 [7] (3+4★) | 6 [6] | | |
| **hiBBB-in-Transwell** | | 33 (TEER only) | −2 | 9 [5] | | 6 [4] | 6 [4] | 5 [4] | 5 [4] |

**Detailed Experimental Section**

Full details on materials and equipment can be found in the last Section.

*Microfluidic Design & Fabrication*

Our system utilizes a vertically-stacked channel geometry common in Barrier-on-Chip systems.[1] A track-etched, 9 μm thick polycarbonate (PC) membrane (1 μm diameter pores, $2\times10^{6}$ $cm^{-2}$) serves as the cell culture support between the channels. The microfluidic channels are fully mirror-symmetric, with a height of 250 μm and an overlap area of 200×15 mm². The channel sidewalls feature a gradual curvature at the membrane interface to aid formation of a continuous cell layer across the interface.

We designed a positive master mold for the channel structures that was CNC-milled in aluminum. We then coated the mold with ~1 μm Parylene to serve as a durable non-stick coating.[2] From this, we fabricated negative PDMS molds on glass (channel tops) or on a block of aluminum (with two drilled holes for later OSTE+ injection; channel bottoms) using standard solution casting methods. Briefly, a 1:10 mixture of PDMS curing agent to base was poured onto the master mold, degassed in a vacuum desiccator, and brought into contact with the oxygen plasma-treated substrate. We applied pressure to remove excess PDMS and cured the weighted-down mold/substrate stacks at 80°C initially on a hotplate (30 min) and subsequently in an oven (1 h). After cooling, we carefully removed the PDMS molds from the master mold.

We followed our previously-published protocol[3] for electrode preparation using laser-cut paper shadow masks. We employed this to pattern interdigitated electrodes (200 nm gold, with 15 nm titanium adhesion layer; 0.3 mm width and 1.5 mm gaps) on the PC membranes. Before use, the electrodes were cleaned by ultrasonicating the membrane in isopropanol (1 min) and then submerging it in deionized water.

The membrane was placed on top of the glass-PDMS top layer mold and aligned to the channel structures. We then aligned the glass-PDMS-membrane stack with the bottom layer aluminum-PDMS mold, and clamped the entire assembly with paper clips. The entire alignment procedure was conducted on an 80 °C hotplate to impart tensile stress on the membrane at room temperature (due to thermal expansion/contraction).

We thoroughly mixed Component A and B of OSTE+ at a ratio of 1.09:1 in a speed mixer, after which the mixture was placed in a vacuum desiccator for at least one hour to eliminate bubbles, and finally placed in the fridge to cool. We loaded this into a plastic syringe, and slowly injected the mixture into the mold assemblies using one of the drilled holes in the aluminum block serving as the bottom part of the mold, taking care to flush out any air bubbles. The mold with injected OSTE+ was then stored at 5 °C for 10 minutes, before we exposed it with a UV lamp (~10 mW/cm² @ 365 nm) in 30 s intervals (with >30 s rest intervals) for 5 min total. The pre-cooling, rest intervals, as well as the use of a Peltier cooler during exposure serve to prevent premature initiation of the second curing step.[4] We carefully removed the semi-cured OSTE+ channel layer with the in-molded membrane from the mold assembly, and diced the layer into individual devices with scissors, subsequently placing them on a transfer support made of TOPAS. We then pressed the layer onto 125 μm thick PC foil (under lateral compressive stress, to provide the membrane with tensile stress upon relaxation) serving as the device bottom. Using needles as guides, the PC-OSTE stack was then aligned and pressed onto the microfluidic connection assembly (CNC-diced from commercial PC parts). These semibonded devices were clamped between a metal holder and glass slides using paper clips, and cured for five days at 75 °C, after which the devices were stored dry at room temperature until usage.

*Maintenance of hiPSC and lt-NES*

We cultured hiPSC in mTeSR 1 in Matrigel®-coated 6-well plates. The cells were passages when reaching 60-80% confluency by first washing one time with Versene solution and then incubated with Versene solution for 4 min at 37°C. We then collected and replated the cells in mTeSR 1. We kept the cells in Essential 8 media for at least two passages before starting a differentiation.

We maintained the neuroepithelial stem cells (lt-NES)[5] in a T25 culture flask that was pre-coated for at least 2h each with 1:500 PLO and 1:500 L2020 (both in DPBS +/+). We cultured the cells in DMEM/F12+Glutamax containing 1% N2 and 0.1% B27 supplement, bFGF and hEGF. The cells were passaged at 80% confluency by incubation with TrypLE for 4 min at 37°C. We then collected the cells in DTI for centrifugation and plated them in the media just described.

All cell cultures were kept in an incubator at 37°C with 5% $CO_2$ and 90% humidity.

*hiPSC differentiation to hiBMEC-like cells*

We followed the protocol by Neal *et al.*[6] In brief, we seeded hiPSC in 6 well plates coated with Matrigel at a density of 15 800 cells/cm$^2$ in E8 medium with 10 µM Y-27632 (Rho kinase inhibitor). After 24h of culture (day 0), we changed the medium to E6 medium to initiate differentiation. We replenished the media daily until day 4, when we switched to human endothelial serum-free media (heSFM) containing 20 ng/ml basic fibroblast growth factor (bFGF), 10 µM retinoic acid (RA), and 2% B-27 and left to culture for another 2 days. On day 6, we washed the cells with DPBS +/+ and incubated with TrypLE for 4 min at 37 °C for singularization. We collected the cells in heSFM and, after centrifugation, transferred them to heSFM supplemented with 10% dimethyl sulfoxide and 30% fetal bovine serum. The cells were frozen by first utilizing a Mr. Frosty freezing container in a –80°C freezer before we transferred them to liquid nitrogen storage. We want to point out that the endothelial identity of these cells has been discussed by us and others.[7–9]

*lt-NES differentiation to hiAstro-like cells*

To initiate lt-NES differentiation (day −1), we split the cells and seeded them in attachment factor (AF)-coated 6-wells at a density of 288 000 cells/well in AM media supplemented with the included FBS and AGS supplements. Media was changed every 48h and the cells were split and reseeded at confluency. On day 28 of the differentiation, the cells were collected in AM supplemented with 10% fetal bovine serum and 10% dimethyl sulfoxide and frozen by first utilizing a Mr. Frosty freezing container in a –80 °C freezer before we transferred them to liquid nitrogen storage. A manuscript with the protocols' biological characterization is forthcoming.[10]

*Transwell culture of hiPSC derived neurovascular unit*

We pre-coated 24 well plate Transwell inserts with a 100 µg/ml : 400 µg/ml fibronectin/collagen IV solution (in DI water) and incubated them at least 4 hours in 37°C up to over-night. We thawed the hiBMEC in a 37° water bath, then slowly added fresh heSFM.

After centrifugation, we transferred the cells to heSFM supplemented with 0.2% Primocin, 2% B-27, 0.1% Y-27632, 10 μM RA, 20 ng/ml bFGF, and 0.01 mg/ml BioSilk, and performed a count. The cells were seeded on to the Transwell insert membranes at $5\times10^5$ $cm^{-2}$.

We precoated 24 wells with 5% L521 in PBS at least 4 hours at 37 °C up to over-night. We thawed hiAstro a 37° water bath, then slowly added fresh AM. After centrifugation, we transferred the cells to AM supplemented with the included FBS and AM supplements, as well as 0.2% primocin and 0.25% L521. We seeded hiAstros at a density 35 000 cells/$cm^2$.

After 24h of separate subculture, the media was changed to heSFM with 0.2% primocin, 1% B-27 (minus Antioxidants) and 0.5% ITS, and hiBMEC-culture Transwell inserts added to hiAstro-containing wells. At 48h, we transitioned the culture to heSFM with 0.2% primocin and 1% ITS.

*Microfluidic culture preparations*

On day −2 we exposed the devices as well as all tubing to oxygen plasma for ~1 minute. We then connected the devices to a 16-channel peristaltic pump (0.25 mm inner diameter tubing) using 0.51 mm inner diameter tubing and 23 g metal pins. Devices were further placed on CNC-machined acrylic platform, where we employed spring-loaded pins to make electrical connection to the device contact pads. Coaxial cables connected this assembly to a relay multiplexer, itself connected to an LCR meter (further details in the CESR measurement section).

We disinfected the devices by flowing 70% ethanol for 5 min after which we rinsed with DPBS +/+ for 10 min. At every step we checked for bubbles and flushed them out as needed. Media reservoirs (5 ml plunger-less syringes with 16 g blunt needles) were then connected via 1.6 mm inner diameter tubing and put inside an incubator. Flow was turned on and monitored with a LabView program over night and media was collected from outlets to ensure equal flow volumes. Our flow scheme consisted of 45 s flow at ~0.45 ml/h, followed by a 225 s pause, yielding an average flow rate of ~1.5 μl/min. Every 6 hours, the system was flushed at 75 μl/min for 10 s to remove potential channel blockages (bubbles, clusters of cell debris).

On day −1 we introduced coatings into the devices. We aspirated 100 μg/ml : 400 μg/ml fibronectin/collagen in the vascular channels and left it to incubate at least 6 hours inside the incubator. Subsequently, we aspirated 50% L521 in the perivascular channels and left it to incubate undisturbed at least 2 hours in the incubator. We subsequently re-connected the media reservoirs, and (after another 6-hour incubation) began perfusing devices with B27-supplemented heSFM.

*Microfluidic culture of hiPSC derived neurovascular unit*

On day 0 we first thawed the hiAstro-like cell and prepared a suspension of 2.25 million cells/ml in AM with FBS and AGS supplement, as well as 2.5% L521 and 0.2% primocin. The cell suspension was aspirated in the perivascular channel at 5 μl/min. The devices were kept "upside down" with the perivascular channel facing up and incubated for 1h in static conditions at 37°C to let the cells attach. We then turned on a slow, constant media flow (~0.3 μl/min) for another 2h. Subsequently, we thawed the hiBMEC-like cells and aspirated them into vascular compartments as a suspension of 50 million cells/ml at 5 μl/min. We utilized heSFM supplemented with 0.2% Primocin, 2% B-27, 0.1% Y-27632, 10 μM RA, 20 ng/ml bFGF, and 0.1 mg/ml BioSilk. The cells were left to attach for 60 min under static conditions at 37 °C, with devices variously angled at ~20 min intervals to ensure even coverage during attachment.

Flow was then turned on at a constant 0.15 µl/min, and over 6 h gradually increased to 1.5 µl/min. On day 1 (24h), of subculture, the media was changed to heSFM with 0.2% primocin, 1% B-27 (minus Antioxidants) and 0.5% ITS. On day 2 (48h), we transitioned the culture to heSFM with 0.2% primocin and 1% ITS.

*Challenging of the barrier*

On day 3, we refreshed the media in both systems (same formulation as d2). The antioxidative reagent NACA was spiked into randomly-selected Transwell vascular compartments and device vascular channel inlets on day 3 (70.5 h) at a final concentration of 4 mM. Nitrosative stress was induced on day 3 (71 h) by spiking 1 mM or 4 mM SIN-1 into the Transwell perivascular compartments or 4 mM into the device perivascular channel inlets. Transwell plates were mounted on a rocking plate inside the incubator to ensure within-compartment mixing.

*Transendothelial electrical resistance measurements*

We measured Transwell TEER (using an EVOM2 with STX3 electrodes) at 48 h, 70 h and 95 h after seeding to monitor tight barrier formation and the effect of the treatment on the barrier tightness. We correct for the resistance of the porous membrane (area $A$ = 0.33 mm²) using coating-only "blank" Transwells kept under the same conditions as the other wells with the equation:

$$\text{TEER}\ [\Omega\ \text{cm}^2] = (\ \text{TEER}[\Omega] - \text{TEER}_{\text{blank}}[\Omega]\ ) \times A[\text{cm}^2]$$

*CESR measurements & analysis*

We connected the LCR meter to the relay multiplexer with coaxial cables, as well as an in-line high-accuracy 5 kΩ resistor. Combined with a 50 mV signal level, this served to limit the maximum current inside the devices (and thus experienced by the cells) to 10 µA. The relay multiplexer distributed the signal (and readout) serially across our up to eight devices per experiment. A LabView UI was written to control measurement timing and data collection. We set it to conduct one impedance readout per device, followed by a 1-minute interval with open relay circuits (5 minutes for d0-d3). Non-zero switching and readout times add an additional 30 seconds to the timings.

After data collection, we proceeded to employ multiple stages of background correction to the raw impedance data. First, we measured open, short, and load (10 nF capacitor, chosen for its broadly similar frequency response with our devices) reference signals the same coaxial cables and spring-loaded contacts as for our devices, and applied the standard correction for these. Second, we subtracted the coating-only impedance measured from each device immediately before cell seeding. Where this correction yielded obvious outliers (due to bubbles in the channels at the correction timepoint; $n$=4), we adjusted toward the average signal (from other devices within that same experimental repeat) at ~3 h post-seeding, where a distinctive dip in signals marks the beginning of the liquid flow ramp. The real part of the resulting impedance signals at 6 kHz yielded the raw CESR values. For normalization during the intervention phase, we further divided each signal trace by the averaged CESR from 3 h prior to the introduction of SIN-1. We employed MATLAB and OriginLab for the data analysis.

*Tracer dye permeability*

To test the permeability of the cell barrier we added 100 μg/ml Cascade Blue hydrazide in the Transwell vascular compartment (70 h) or device vascular channel inlet (48 h and 70 h). We collected samples at 70 h (devices only) and endpoint, and prepared serial dilutions from the relevant inlet solutions as reference. We then used a fluorescence plate reader and measured signal *F* at $\lambda_{ex/em}$ = 400 / 430 nm. To determine (fractional) concentrations *C*, the calibration samples were fitted to the empirical model (chosen based on simplicity and goodness-of-fit):

$$\log C[1] = a \cdot (\log F)^b + c$$

Due to the wide range of concentrations, we used separate models for the low- and high-concentration ranges (using a cutoff of *C*~4%, chosen to yield a smooth model transition).

With the actual samples, we then used the following equation to calculate the permeability of the devices:

$$P'_{\text{app}}[\text{cm s}^{-1}] = \frac{C_{\text{out}}^{\text{perivasc.}}[1] \cdot Q[\text{cm}^3\ \text{s}^{-1}]}{A[\text{cm}^2]}$$

where $Q$ is the average volumetric flow rate and $A$ the permeable membrane area.

For Transwells, the formula changes due to accumulation of dye over time *t*=24 h inside the perivascular compartment with volume *V* = 800 μl.

$$P'_{app}[\text{cm s}^{-1}] = \frac{V^{\text{perivasc.}}[\text{cm}^3] \cdot C_{\text{out}}^{\text{perivasc.}}[1]}{A[cm^2] \cdot t[s]}$$

Additionally, we correct for the resistance of the membrane with samples from “blank” (coating-only) measurements, taken either at d−1 (devices) or in parallel with other permeability measurements (Transwells):

$$P_{\text{app}} = \left(\frac{1}{P'_{\text{app}}} - \frac{1}{P_{\text{app}}^{\text{blank}}}\right)^{-1}$$

*Immunocytochemistry*

The cell media was removed and Transwells and devices were washed with DPBS +/+ before fixation with 10 % formalin solution for 20 min at RT. Device washes (and other solution exchanges) were performed by pump perfusion at ~ 5 μl/min, whereas Transwells were pipetted. After three washes with DBPS we added blocking buffer consisting of 10 % goat serum and 0.1% Triton-X in DPBS −/− for 1h at RT. We prepared antibody solutions and nuclear stain solution in a dilution buffer consisting of DPBS −/− supplemented with 1% goat serum and 0.01 % Triton-X. After removing the blocking buffer, we added primary antibody solution containing 7.5 μg/ml anti-ZO-1 and 20 μg/ml anti-S100B in devices. In Transwell vascular compartments we added 3.75 μg/ml anti-ZO-1 and in the Transwell perivascular compartments 10 μg/ml anti-S100B (subset: additionally 2 μg/ml anti-CD44). All primary antibody solutions were incubated over night at 4 °C. After incubation we removed the primary antibody solutions and washed three times with 0.01 Triton-X in DPBS −/−. We then added secondary antibody solution containing 4 μg/ml anti-mouse IgG and 4 μg/ml anti-rabbit IgG in devices. In Transwell apical compartments we added secondary antibody solution containing 2 μg/ml anti-mouse IgG, and in Transwell basal compartments we added 2 μg/ml anti-rabbit IgG (subset: additionally 2 μg/ml anti-mouse IgG) solution. The secondary antibody solutions were left to incubate for 1h in RT protected from light after which we removed the solution and washed twice with 0.01% Triton-X DPBS −/−. Both devices and Transwells were then counterstained with a solution of 5 µg/ml Hoechst by 10 min incubation at RT protected from light. We then removed the Hoechst solution and washed three times with 0.01% Triton-X DPBS −/−. Devices were filled with VectaShield mounting medium and stored in darkness at 4 °C until imaging. Transwell membranes were cut out from the apical compartment with a scalpel and mounted with VectaShield between a glass slide and glass coverslip. From the basal compartment of the Transwells, we removed the DPBS solution and added VectaShield mounting medium under a glass coverslip. Devices and transwells were imaged with a laser scanning microscope. Widefield images of Transwell bottom wells were taken with widefield microscope.

*Image analysis*

All image analysis was carried out using Fiji.[11] Confocal images were assembled using maximum projection and brightness/contrast was adjusted for inadvertent differences in staining intensities between experimental repeats/groups to emphasize morphology. Quantification of hiAstro surface coverage in Transwells was done using the Li threshold algorithm, and then measuring the green signal fraction across the entire cell growth area. 1% coverage was assigned manually to images with insufficient cell/background contrast, based on visual comparisons with successfully quantified low-coverage wells. High-magnification hiBMEC images were thresholded using the Huang algorithm, and analyzed using the Particle Analysis module.

*UPLC/MS metabolomic analysis & processing*

For devices, we collected outflow samples over pre-intervention (d3) and at endpoint (d4). For Transwells, we collected samples only at endpoint (d4). Sample numbers are given in Table S4. Samples were frozen down immediately after fluorescence measurement (cf. permeability assays), stored at −80 °C, and shipped on dry ice. Additionally, $N$ = 14 media control samples from device inlets or the no-cell Transwell were included.

A detailed description of the technical setup is available in a prior publication.[12] In brief, we utilized a dual-polarity LC-MS on an Orbitrap ID-X coupled to a ZIC-pHILIC column. Quality control samples were prepared from pooling assay sample aliquots, and run every 11 samples. Pooled sample aliquots, grouped by compartment (TW/device/vascular/perivascular), were also utilized for acquiring MS/MS-level data.

Analysis pipeline 1, aimed at individual compounds, also followed the aforementioned paper, relying on vendor software. Briefly, Compound Discoverer was used for untargeted sample analysis using the corresponding built-in workflow, relying on both MS1 and MS2-level data. This pipeline yielded ~2700 compounds, of which 161 could be assigned tentative annotations. For top-scoring compounds, annotation likelihood is based on MZcloud library match and FiSH scoring. To classify a compound as endogenous or not, we relied on HMDB.[13] Features corresponding to NACA and SIN-1C were further integrated in TraceFinder for our semi-quantitative/qualitative concentration analysis.

Analysis pipeline 2, aimed at network-level data, relies on open-source tools and is adapted from another of our prior metabolomic studies.[14] We converted raw data using ProteoWizard,[15] and relied on XCMS for feature alignment and grouping.[16] Parameters were derived from guided optimization (on quality control samples) with algorithms from the AutoTuner,[17] patRoon,[18] and MetaoAnalystR[19] packages. The data and XCMS analysis parameters are available online. This pipeline yielded ∼3700 features after applying filters for mass (65–2000 Da), retention time (excluding leading/trailing peaks of the injection), and signal-over-blank ratio (>6). Remaining missing values in the feature table – which at this stage are present only where raw signals are below the detection limit – were filled using a QRILC algorithm.[20] Drift compensation was performed with the statTarget package based on the quality control samples (QC-RFSC algorithm).[21] We further discarded features with post-correction QC RSD > 40%, leaving us with ~3300 features for network analysis.

Data from both pipelines were then adjusted for media background by subtracting the median of the corresponding media sample group (SIN-1, NACA, neither; or, particularly with broken barriers, appropriate linear combinations thereof) from a given assay sample. To minimize the number of negative signals in subsequent analysis, we then added back in – to all samples/features – the highest median signal of the three media groups. Data were then fed into statTarget for multivariate and univariate analysis, employing a generalized-log2 transformation and mean-centering.

We then took the *p*-values and glog2-changes from pipeline 2 for network analysis using the mummichog + GSEA approach available in MetaboAnalystR. Based on pipeline 1 results, we chose a mass accuracy cutoff of 1 ppm. We chose the p-value in line with a suggested target of 10% hits, an average we came closest to with $p = 0.01$. As noted in our prior publication, the combined pathway analysis is unstable upon repeat analysis. Therefore, we combined *p*-values and NES scores from 25 analysis runs by geometric[22] and arithmetic averaging, respectively, before integrating the respective mummichog and GSEA results through Fisher's method as intended by the authors.[23]

### Materials & Equipment

#### *Cells*

We used the hiPSC line CTRL-9-II described by Uhlin et al.[24] as provided by the iPS Core Facility at Karolinska Institute under a Material Transfer Agreement. The iPS core further provided us with corresponding lt-NES from the same line, derived according to the protocol by Falk et al.[5]

#### *Immunocytochemistry*

| Product | Manufacturer | Identifier |
|---|---|---|
| Formalin solution, neutral buffered, 10% | Sigma-Aldrich | HT501128 |
| Anti-ZO-1 Monoclonal Antibody (IgG1, Mouse) | Thermo Fisher | 33-9100 |
| Anti-S100 beta Monoclonal Antibody (IgG, Rabbit) | Abcam | ab52642 |
| Anti-CD44 Monoclonal Antibody (IgG2a, Mouse) | Thermo Fisher | MA-13890 |
| Anti-Mouse IgG CF 568 antibody | Sigma-Aldrich | SAB4600083 |
| Anti-Rabbit IgG CF 488A antibody | Sigma-Aldrich | SAB4600044 |
| Hoechst 33342 | Gibco | H3570 |
| Goat Serum | Sigma-Aldrich | G9023 |
| Triton X-100 (1%) | Gibco | HFH10 |
| Vectashield Antifade mounting medium | Vector Labs | H-1000 |
| Cascade Blue hydrazide, trisodium salt | Thermo Fisher | C687 |

### *Cell Culture*

| Product | Manufacturer | Identifier |
|---|---|---|
| mTeSR 1 feeder-free maintenance medium | STEMCELL Technologies | 85857 |
| Matrigel Growth Factor Reduced Basement Membrane Matrix | Corning | 354230 |
| Versene Solution | Gibco | 15040066 |
| Essential 8 (E8) Medium | Gibco | A1517001 |
| Human Endothelial Serum Free Medium | Gibco | 11111044 |
| Recombinant Human FGF basic/FGF2/bFGF | RnD Systems | 233-FB |
| Y-27632 dihydrochloride | RnDSystems | 1254 |
| Retinoic Acid | Sigma Aldrich | R2625 |
| B-27 serum free supplement (50X) | Gibco | 17504044 |
| TrypLE Select Enzyme (1X) | Gibco | 12563029 |
| DPBS, + calcium, + magnesium | Gibco | 14040091 |
| DPBS, − calcium, − magnesium | Gibco | 14190144 |
| Dimethyl sulfoxide | Sigma Aldrich | D2650 |
| Fetal Bovine Serum (One Shot) | Gibco | A3160801 |
| Primocin | Invivogen | ant-pm |
| Fibronectin from bovine plasma | Sigma Aldrich | F1141 |
| Collagen IV from human placenta | Sigma Aldrich | C5533 |
| Recombinant human laminin 521 (L521) | BioLamina | LN521 |
| Astrocyte medium (AM) | ScienCell | 1801 |
| Fetal Bovine Serum (FBS) | ScienCell | 0010 |
| Astrocyte Growth Supplement (AGS) | ScienCell | 1852 |
| DMEM/F-12, GlutaMAX | Gibco | 31331028 |
| N-2 Supplement | Gibco | A1370701 |
| Recombinant spider silk protein (Biosilk) | BioLamina/Spiber | BIOSILK |
| Poly-L-ornithine hydrobromide | Sigma Aldrich | 3655 |
| Laminin from EHS murine sarcoma basement membrane (L2020) | Sigma Aldrich | L2020 |
| Recombinant human epidermal growth factor (EGF) | Sigma Aldrich | E9644 |
| Defined Trypsin Inhibitor | Gibco | R007100 |
| Attachment factor protein (1X) | Gibco | S006100 |
| *N*-Acetylcysteine amide | Tocris | 5619 |
| SIN-1 chloride | Biotium | 00221-1 |

### *Consumables & Microfluidics*

| Product | Manufacturer | Identifier |
|---|---|---|
| 6-well plate, tissue culture-treated | VWR | 734-2323 |
| 24-well plate, tissue culture-treated | Corning | 3526 |
| Transwell Permeable Supports (6.5mm, 0.4 µm pores, polyester) | Corning | 3470 |
| 96-well half area plate | Corning | 3882 |
| Microfluidic interface (polycarbonate, microscopy slide format, 2x16 olives) | Microfluidic chipshop | 10-1121-0343-03 |
| Polycarbonate film (125 µm) | Covestro | Makrofol DE 1-1 |
| Syringes, 5ml (as liquid reservoirs) | Restek | 22774 |
| Blunt dispensing needles 16G | Metcal | 915100-TE |
| Steel Microfluidic Fittings 23G | Elveflow | LVF-KFI-13 |

| | | |
|---|---|---|
| PharMed peristaltic pump tubing (0.25mm ID) | Ismatec | SC0320 |
| Pharmed extension tubing (0.51mm ID) | Ismatec | SC0339 |
| Tygon ND 100-65 Medical Tubing (1/16" ID) | Saint-Gobain | ADF00002 |
| ipCELLCULTURE track-etched porous membrane (10µm thick, 1 µm pores, 2E6 $cm^{-2}$) | It4ip | (N/A) |
| OSTE+ Crystal Clear | Mercene Labs | OSTEMER 322 |

*Equipment*

| Product | Manufacturer | Identifier |
|---|---|---|
| $CO_2$ incubator | Thermo Scientific | Heracell Vios 160i |
| Mr. Frosty Freezing Container | Gibco | 5100-0001 |
| 16-channel peristaltic pump with click-'n-go cartridges | Ismatec | IPC-N |
| Switch Control Unit and 10 Channel Relay Multiplexer Card | HP/Agilent | 3488A with 44470A |
| RCL (LCR) Meter | Fluke/Philips | PM 6304 |
| Epithelial Volt/Ohm (TEER) Meter | WPI | EVOM2 |
| Chopstick Electrode Set | WPI | STX3 |
| Multimode Plate Reader | Tecan | Infinite 200 Pro |
| Confocal Laser Scanning Microscope | Zeiss | LSM800 |
| Widefield Microscope | Olympus | IX73 |
| Orbitrap ID-X LC-MS (Q-Exactive plus in line with Ultimate 3000 LC) | Thermo Fisher | (N/A) |
| ZIC-pHILIC column (2.1×150 mm, 5 µm) | Millipore | 150460 |
